# Hippocampal-to-Ventricle Ratio as a Head-Size-Independent Biomarker: Sex Differences and Cognitive Associations in 27,680 UK Biobank Participants

**DOI:** 10.64898/2026.04.13.718285

**Authors:** Sofia Fernandez-Lozano, D. Louis Collins

## Abstract

Hippocampal volume is a key biomarker for Alzheimer’s disease and brain aging, yet reported sex differences vary substantially depending on head-size adjustment methods. We examined whether the hippocampal-to-ventricle ratio (HVR), a self-normalizing measure, provides more consistent sex difference estimates than conventional adjustment approaches. Using a subset of UK Biobank structural MRI data (N = 27,680; 56% female; ages 45–82), we compared sex differences across four adjustment methods (unadjusted, proportions, stereotaxic, residualized) and in samples matched on age and on intracranial volume (ICV). Hippocampal sex differences reversed direction across methods, ranging from *d* = −0.89 (males larger, unadjusted) to *d* = 0.58 (females larger, proportions), with a range of 1.5 standard deviations across analytical choices. In contrast, HVR showed a consistent female advantage (*d* = 0.52) that persisted in the ICV-matched subsample (*d* = 0.19), confirming this effect is not a head-size artifact. Males exhibited steeper cross-sectional age-related HVR differences (1.7× female rate, *p* < 10^−50^), consistent with males showing ventricular expansion at younger ages. Structural equation modeling revealed that HVR predicted general cognition comparably to hippocampal volume (*β* = 0.04, overlapping CIs), and unlike residualized hippocampal volume, HVR maintained significant brain-cognition associations in both sexes. We provide sex-specific normative centile curves for clinical application. These findings indicate that apparent hippocampal sex differences largely reflect methodological choices rather than biology, while HVR captures consistent morphological variation related to brain aging that may have clinical utility.

## 1 Introduction

The hippocampus (HC) is a brain structure used as an early biomarker for Alzheimer’s disease (AD) and brain aging [Jack et al., 1999; Jack et al., 2010; Raz et al., 2005]. Sex differences in brain structures have been documented, but their interpretation is controversial [Ritchie et al., 2018; Ruigrok et al., 2014], largely because findings depend on whether and how head-size adjustments are applied [Nordenskjöld et al., 2015; Sanchis-Segura et al., 2020]. Males have approximately 12% larger intracranial volume (ICV) than females [Ruigrok et al., 2014], and after head-size correction, sex classification from brain features drops from >80% to approximately 60% [Sanchis-Segura et al., 2020]. This methodological sensitivity suggests some reported sex differences may be artifacts of adjustment methods.

Hippocampal atrophy rates predict conversion from mild cognitive impairment to dementia [Rajagopal et al., 2024], making accurate volume assessment clinically consequential. However, males and females show different rates of age-related brain volume change, with steeper subcortical volume declines in males across multiple structures [Ritchie et al., 2018]. These sex-differential trajectories mean that normative reference values must be sex-stratified to avoid misclassifying normal variation as pathological; recent large-scale brain charting efforts have adopted sex-specific centile modeling for this reason [Bethlehem et al., 2022; Ge et al., 2024].

Multiple adjustment methods exist, each with different assumptions: proportions (V/ICV) assumes perfect allometric scaling, residualization (V − *β*× ICV) removes linear ICV association, and stereotaxic normalization applies spatial transformation to a template. These methods can yield contradictory conclusions about the same brain regions [Wang et al., 2024]. For hippocampus, reported sex differences range from males larger (unadjusted; Ruigrok et al. [2014]) to females larger (proportions; Sanchis-Segura et al. [2020]) to no difference (residualized; O’Brien et al. [2011]). The proportions method can produce biologically implausible results, while residualization yields more consistent estimates [Wang et al., 2024]. No consensus exists on the correct method, and conclusions depend on analytical choices.

The hippocampal-to-ventricle ratio (HVR), defined as HC / (HC + LV) where HC denotes total hippocampal volume and LV denotes the volume of the temporal horns of the lateral ventricles, offers a potential solution. Both structures scale with head size, making their ratio self-normalizing. HVR captures the relationship between hippocampal and ventricular volumes, which show opposite patterns with age—smaller hippocampus and larger ventricles. Schoemaker and colleagues introduced HVR and showed that it provides stronger associations with age and memory than standard hippocampal volume in healthy aging [Schoemaker et al., 2019]. Subsequent work shows HVR provides higher effect sizes for distinguishing AD from controls compared to HC alone [Fernandez-Lozano et al., 2025]. However, properties of HVR in healthy populations and across adjustment methods remain unexplored.

Age-related ventricular expansion reflects concurrent loss of surrounding gray and white matter tissue [Madsen et al., 2015], and this expansion occurs earlier in males, with abnormal enlargement appearing in the seventies and preferentially affecting the frontal horns [Currà et al., 2019]. These sex-differential ventricular trajectories suggest that a ratio incorporating both hippocampal and ventricular volumes may capture sex differences in brain aging that individual volumes miss. Furthermore, brain structure–cognition associations themselves may differ by sex: grey matter volumes predict cognitive profiles with different strength and spatial specificity in older males versus females [Jockwitz et al., 2024], and hippocampal volume predicts associative memory in older women but not men [Zheng et al., 2017]. Whether HVR shows similar sex-differential cognitive associations remains unknown.

We addressed three questions: (1) Do hippocampal sex differences vary across head-size adjustment methods while HVR remains consistent? (2) Do cross-sectional age patterns differ by sex, and do these patterns differ between HC and HVR depending on normalization method? (3) Does HVR show comparable cognitive associations to conventionally adjusted hippocampal volumes?

## 2 Materials and Methods

### 2.1 Data Source and Participants

Data were obtained from UK Biobank (Application 45551) [Miller et al., 2016; Sudlow et al., 2015]. The initial imaging sample comprised approximately 47,398 participants with T1-weighted structural MRI. After exclusions (detailed below), the final analytic sample included N = 27,680 participants aged 44.6 to 82.8 years. With this sample size, we had >99% power to detect small effects (d = 0.10) and >80% power to detect very small effects (d = 0.05) for two-group comparisons at *α* = .05.

Three additional subsamples were defined for specific analyses (Figure 1; Table 1). The **England subsample** (N = 25,172) comprised participants assessed at UK Biobank centres in England and was used for SEM analyses requiring the Index of Multiple Deprivation (IMDP), which is not comparable across UK countries.

**Figure 1:**
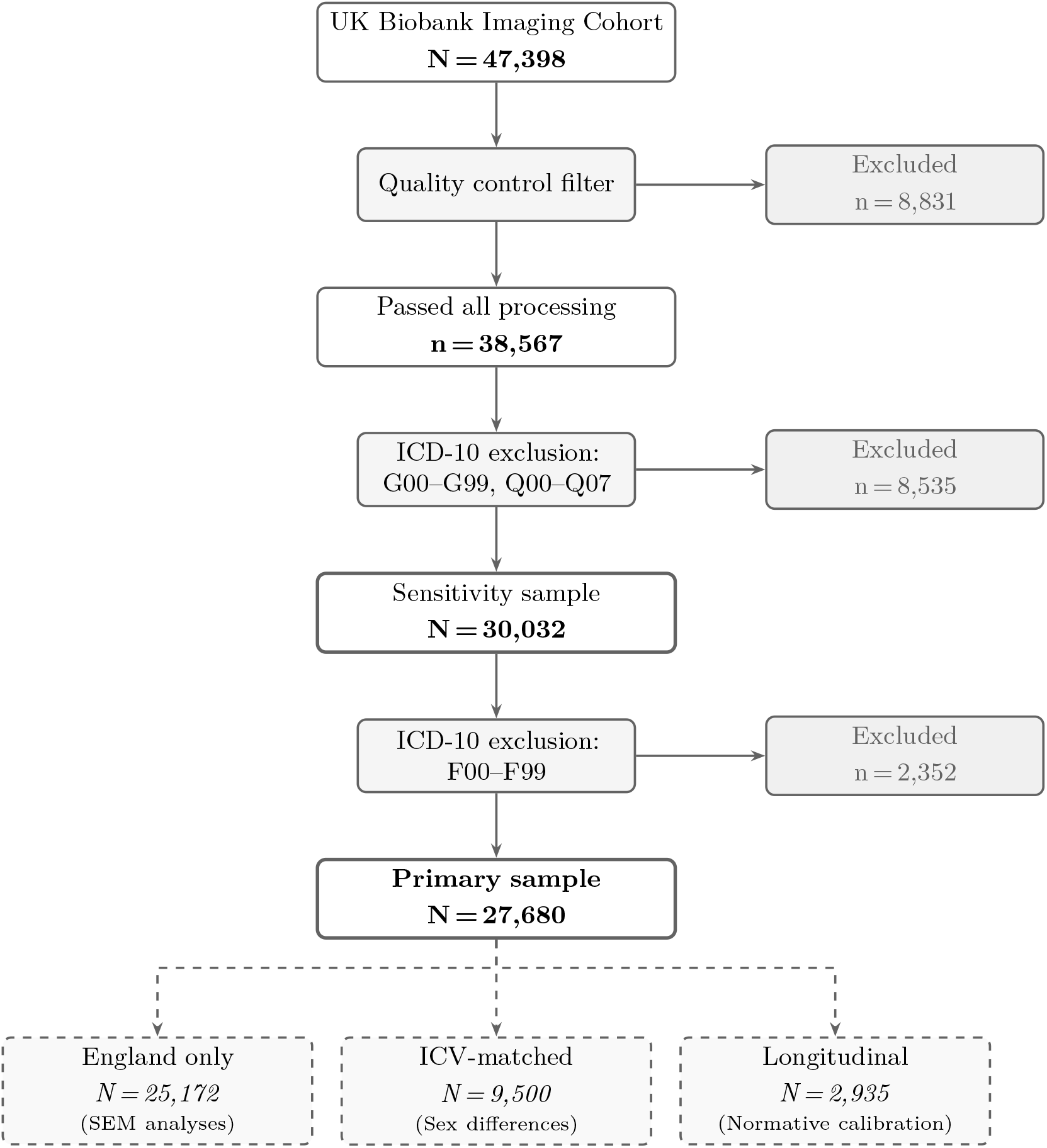
Participant selection flow. Dashed boxes indicate analytic subsamples derived from the primary sample.

**Table 1:**
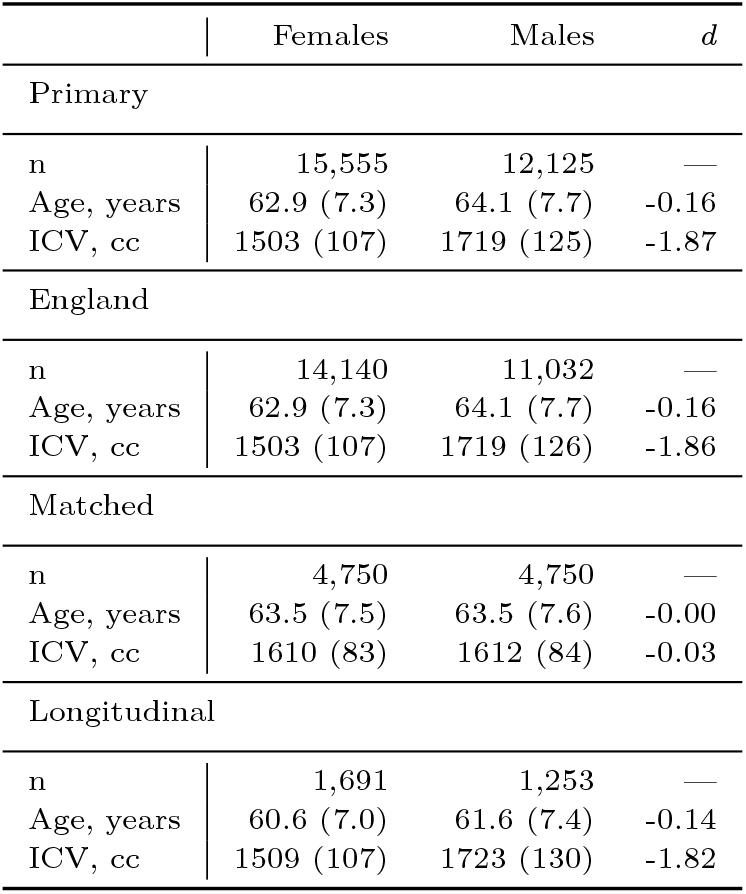
Participant characteristics by sex and sample. Values are M (SD). d = Cohen’s d (Female − Male). England = English assessment centres (SEM analyses). Matched = age ±1 yr, ICV ±25 cc. Longitudinal = participants with repeat imaging.

This subsample provides adequate power for detecting standardized path coefficients ≥ 0.02. The **ICV-matched subsample** (N = 9,500) comprised female–male pairs matched on age (±1 year) and ICV (±25 cc) drawn from the primary sample, and was used to isolate sex effects from head-size confounding (see Section 2.4.1). The **longitudinal subsample** (N = 2,935) comprised participants from the primary sample who also had a follow-up imaging session (ses-3), with a mean intersession interval of 2.7 years (SD = 1.1; range: 1.5–5.0). This subsample was used as a sensitivity check for normative model calibration (see Section 2.4.2).

#### 2.1.1 Exclusion Criteria

Participants were excluded if they had any of the following ICD-10 diagnoses recorded in linked hospital records: F00-F99 (mental and behavioral disorders including dementia, mood disorders, schizophrenia, and anxiety disorders, which are associated with hippocampal volume alterations [Brosch et al., 2022; Cao et al., 2023; Schmaal et al., 2016]); G00-G99 (neurological diseases including Parkinson’s disease, multiple sclerosis, epilepsy, and cerebrovascular disease, which directly affect brain structure [Erp et al., 2016; Koenig et al., 2014; Tai et al., 2025; Xu et al., 2020]); and Q00-Q07 (congenital CNS malformations affecting normal brain development). This exclusion strategy selected a sample of cognitively healthy adults without conditions known to affect hippocampal morphometry. The full exclusion flow is shown in Figure 1.

#### 2.1.2 Quality Control

Image segmentations underwent quality control in three stages: (1) automated quality scores from deep learning-based assessments, requiring DARQ score ≥ 0.25 [Fonov et al., 2022] and AssemblyNet rating of “A” (highest quality grade) [Coupé et al., 2020], (2) statistical outlier detection excluding participants with age- and head-size-adjusted volumes exceeding ±3 SD from the sample mean, and (3) visual inspection of flagged cases by trained raters for evident segmentation failures, motion artifacts, or anatomical anomalies. This multi-stage pipeline is intentionally conservative: in a large population-based study where statistical power is not a limiting factor, prioritizing data quality over sample retention reduces the risk of segmentation errors biasing volumetric estimates, particularly for small structures like the hippocampus and temporal horn ventricles where even minor segmentation failures can produce large relative errors.

For participants with multiple imaging sessions, QC filtering was applied first, then one session per participant was selected for cross-sectional analyses. When a participant’s baseline session (ses-2) failed QC but the follow-up session (ses-3) passed, the follow-up session was retained in the cross-sectional sample. Longitudinal analyses required both sessions to pass QC.

### 2.2 Brain Measures

#### 2.2.1 Image Acquisition

Images were acquired on Siemens Skyra 3T scanners at three UK sites (Cheadle, Newcastle, Reading) using a T1-weighted MPRAGE sequence at 1mm isotropic resolution. Full protocol details are provided in Alfaro-Almagro et al. [2018].

#### 2.2.2 Segmentation

Hippocampal and ventricular volumes were segmented using an in-house convolutional neural network trained on manually labeled data. Validation against manual segmentations showed Dice *κ* = 0.94 and Pearson r = 0.99 [Fernandez-Lozano et al., 2025]. Outputs included bilateral hippocampal volumes (left, right, total) and temporal horn volumes of the lateral ventricles (left, right, total). Intracranial volume (ICV) was obtained from AssemblyNet [Coupé et al., 2020], a large-ensemble convolutional neural network for whole-brain MRI segmentation.

#### 2.2.3 Head-Size Adjustment Methods

Four adjustment approaches were applied to hippocampal and ventricular volumes [Sanchis-Segura et al., 2020; Sanfilipo et al., 2004]:

1. **Unadjusted**: raw volumes in native space (mL), which preserve natural variation but are confounded by overall head size.
2. **Proportions**: volume expressed as a proportion of ICV, which assumes isometric (1:1) scaling and may over- or under-correct depending on the true allometric relationship [Jong et al., 2017; Williams et al., 2021]:

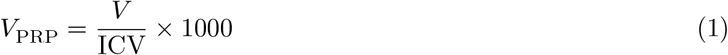
3. **Stereotaxic**: volumes in standard template space after nonlinear registration, which corrects for global size and shape differences:

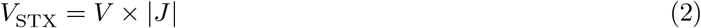

where *J* is the Jacobian determinant of the nonlinear transformation.
4. **Residualized**: linear regression residuals after removing ICV association, with regression coefficients estimated from the pooled sample [O’Brien et al., 2011; Sanchis-Segura et al., 2020]:

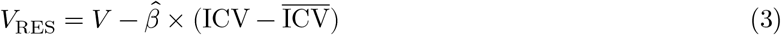

Residualization eliminates ICV-related variance and produces more replicable sex differences compared to proportional methods.

#### 2.2.4 HVR Calculation

The hippocampal-to-ventricle ratio (HVR) was computed as:

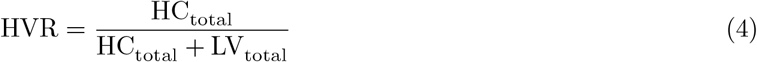

where HC_total_ and LV_total_ denote bilateral sums (left + right) of hippocampal and lateral ventricular temporal horn volumes, respectively. HVR is bounded between 0 and 1, with higher values indicating larger hippocampus relative to ventricles. Because HVR is a ratio of two head-size-dependent structures, no further head-size adjustment was applied.

### 2.3 Cognitive Assessment

#### 2.3.1 Test Battery

Cognitive data were collected at the imaging visit using the UK Biobank touchscreen-based cognitive battery [Sudlow et al., 2015], comprising 12 indicators spanning memory, processing speed, executive function, and reasoning domains (Supporting Information, Table S1). Timed measures (reaction time, trail making, pair matching response time) were inverse-coded so that higher scores consistently indicate better performance across all indicators. Missing cognitive data were handled using full information maximum likelihood (FIML) estimation, which uses all available data under a missing-at-random assumption.

#### 2.3.2 Factor Structure: Bifactor CFA

Cognitive indicators were modeled using a bifactor confirmatory factor analysis (CFA) [Sudlow et al., 2015] with three latent factors. The general factor (*g*) loaded on all 12 indicators. The memory-specific factor (Memory_s_, orthogonal to *g*) loaded on pair matching, numeric memory, and prospective memory. The processing speed-specific factor (Speed_s_, orthogonal to *g*) loaded on reaction time, trail making numeric, and symbol digit substitution correct. Factor scales were set using the marker variable method (first loading fixed to 1.0 for each factor). All latent factors were constrained to be orthogonal, consistent with bifactor model assumptions. Cross-loadings were fixed to zero. Residual covariances were specified for indicator pairs sharing method variance within the same test (pair matching time with errors; symbol digit correct with attempts; trail making numeric with alphanumeric).

Models were estimated using maximum likelihood with robust standard errors (MLR) in lavaan [Rosseel, 2012].

### 2.4 Statistical Analyses

#### 2.4.1 Sex Differences Across Adjustment Methods

##### 2.4.1.1 HVR Self-Normalizing Properties

To evaluate whether HVR reduces head-size confounding, we computed Pearson correlations between ICV and each brain measure (HC, LV, HVR) as well as between ICV and each head-size adjusted HC volume (proportions, stereotaxic, residualized). We expected a self-normalizing measure to show reduced correlation with ICV compared to unadjusted volumes.

##### 2.4.1.2 Sex Differences in Brain Measures

Sex differences were quantified using Cohen’s d with 95% confidence intervals [Cohen, 2013], with positive d indicating females > males by our coding convention. Effect sizes were computed for all ROI × adjustment method combinations in the full sample (N = 27,680).

To isolate sex effects from head-size confounding, we created an ICV-matched subsample where females and males were paired on age (±1 year) and ICV (±25 cc, approximately 0.3 SD). Matching used the MatchIt package (v4.7.2; Ho et al. [2011]) with optimal 1:1 matching without replacement (`method = “optimal”, distance = “mahalanobis”`), with calipers enforcing tolerance bounds. This yielded N = 4,750 matched pairs (total N = 9,500). We reasoned that if sex differences in HC persist after matching on ICV, they cannot be attributed solely to head-size confounding, and if HVR shows similar effects in full and matched samples, this indicates HVR is independent of head size.

#### 2.4.2 Age-Related Patterns and Normative Modeling

##### 2.4.2.1 Age × Sex Interactions

Linear regression models tested whether cross-sectional age-related patterns differed by sex:

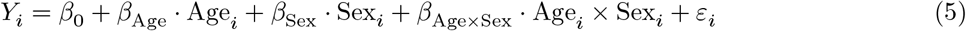

where *Y* is the brain measure, Sex is coded as a binary indicator (Female = reference), and *ε*_*i*_ ∼ 𝒩 (0, *σ*^2^). A significant *β*_Age×Sex_ indicates different age slopes between sexes. Sex-specific slopes were derived as 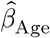for females and 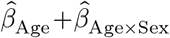 for males, and their ratio quantified the relative rate of cross-sectional age-related differences. Separate models were fit for HC_RES_ and LV_RES_ (both residualized against ICV; Equation 3) and HVR (self-normalizing; Equation 4). Scanner site was not included as a random effect given the low site ICCs (< 0.05; see Sensitivity Analyses), indicating minimal site contribution to variance. While age trajectories may be non-linear (as captured by the GAMLSS normative models below), linear interaction models were used here to provide interpretable summary statistics for sex-differential aging rates.

##### 2.4.2.2 Normative Centile Curves

Sex-specific normative centile curves were estimated using Generalized Additive Models for Location, Scale, and Shape (GAMLSS) [Bethlehem et al., 2022; Rigby and Stasinopoulos, 2005] on the N = 27,434 participants from the primary sample with complete education and site data. For HC and LV (unadjusted):

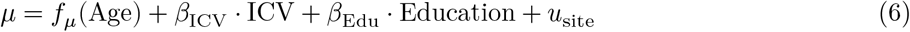

For HVR (self-normalizing, no ICV term needed):

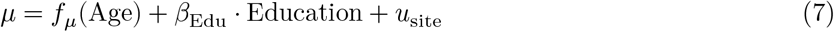

where *f_μ_* (⋅) denotes a cubic spline smooth for the location parameter and *u*_site_ denotes random intercepts for scanner site. Although site ICCs were low (< 0.05; see Sensitivity Analyses), site random intercepts were retained in the normative models because centile estimation targets individual-level classification, where even small systematic shifts can affect percentile assignment. The scale parameter was also modeled as a function of age, *σ* = *f_σ_*(Age), allowing variance to change across the age range. Model families (Gaussian and Box-Cox Cole and Green; BCCG) were compared using AIC, with BIC as secondary criterion. The BCCG distribution was used when flexible modeling of location (*μ*), scale (*σ*), and skewness (*v*) was warranted; shape parameters beyond *σ* were held constant to maintain parsimony. Centile curves (5th, 10th, 25th, 50th, 75th, 90th, 95th) were generated for ages 45–82 by sex.

Data were split into 80% training and 20% test sets, stratified by sex and 5-year age bins, to evaluate model fit on independent data and guard against overfitting. Distribution families were selected on training data and evaluated on the held-out test set by comparing observed vs. expected proportions across centiles. Final models were then fit on the complete cross-sectional sample using the selected distribution family. Model calibration was additionally assessed using the longitudinal subsample (N = 2,935; see Data Source and Participants). Because this subsample comprises the same participants as the primary cross-sectional sample with an additional follow-up session, it tests temporal stability rather than independent validation. Test-retest reliability was quantified using Pearson correlation and intraclass correlation coefficient (ICC, two-way random, absolute agreement).

Two sets of GAMLSS models were generated: (1) site-controlled models including random site intercepts (*u*_site_) for optimal within-UKB accuracy, and (2) transfer models without site effects for application to external cohorts where site is unknown or not comparable. Transfer models used identical distribution families but omitted the random site term, following recommendations for cross-study normative modeling.

#### 2.4.3 Brain-Cognition Associations

Joint measurement-structural equation models tested associations between brain measures and cognitive factors. These analyses were restricted to the England subsample (N = 25,172; see Data Source and Participants) because the Index of Multiple Deprivation (IMDP), used as a socioeconomic covariate, is not comparable across UK countries. All models were estimated in lavaan [Rosseel, 2012].

##### 2.4.3.1 Measurement Invariance

Comparing brain-cognition associations across sexes requires that the cognitive measurement model is equivalent across groups. Measurement invariance of the bifactor structure (see Factor Structure: Bifactor CFA) was therefore tested following standard multi-group CFA procedures [Putnick and Bornstein, 2016]. Four nested models were tested: (1) configural invariance (same factor structure), (2) metric invariance (equal loadings), (3) scalar invariance (equal intercepts), and (4) strict invariance (equal residuals). Invariance was evaluated using ΔCFI ≤ 0.01 [Cheung and Rensvold, 2002] and ΔRMSEA ≤ 0.015 [Chen, 2007]. At minimum, metric invariance is required to compare structural paths across groups.

##### 2.4.3.2 SEM Specifications

Given adequate measurement invariance, three SEM models were estimated, each combining the bifactor cognitive measurement model with structural paths from a brain measure to all three cognitive factors. Brain measures entered as observed (manifest) variables: HC (bilateral hippocampal volume, sum of left and right), HVR (Equation 4), and HC_RES_ (HC residualized against ICV within sex, yielding r ≈ 0 with ICV by construction). The structural component of each model took the following form:

###### Model 1 (HC)

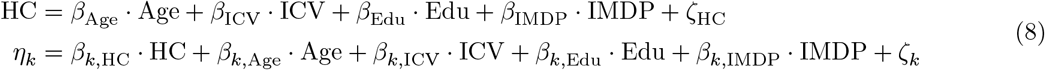

###### Model 2 (HVR)

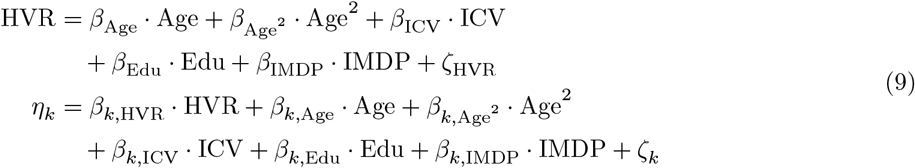

###### Model 3 (HC_RES_)

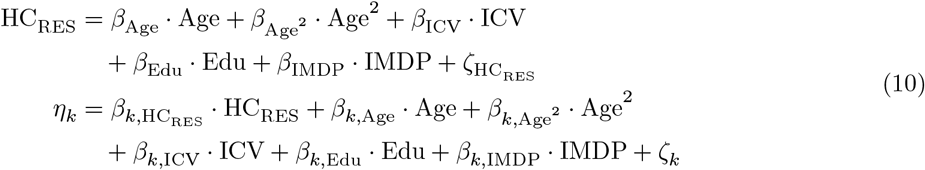

where *η*_*k*_ ∈ {*g*, Memory_*S*_, Speed_*S*_} denotes latent cognitive factors and *ζ* denotes residual terms. A quadratic age term was included in the HVR and HC_RES_ models but not in the HC model, where the combination of raw hippocampal volume, ICV, and quadratic age produced multicollinearity. This difference in model specification means that brain-cognition path comparisons between HC and the other measures should be interpreted with this constraint in mind, as the HC model captures only linear age effects. Age was centered within sex groups before squaring. All structural variables were z-standardized within sex groups to address scale differences. Covariates are summarized in Table S2.

The HC_RES_ model serves as a benchmark representing the methodological standard for removing head-size confounding. Comparing HVR to HC_RES_ tests whether HVR provides similar cognitive associations while being self-normalizing and capturing additional variance through its incorporation of ventricular dynamics. The proportions and stereotaxic adjustment methods were excluded due to concerns about biologically implausible associations and residual ICV correlation, respectively [Wang et al., 2024].

##### 2.4.3.3 Estimation and Inference

Two estimation approaches were used. Pooled models combined all participants, included sex as a covariate, and were estimated using robust maximum likelihood (MLR) with cluster-robust standard errors to account for scanner site. Multi-group models estimated separate structural parameters for females and males, allowing sex differences in brain-cognition paths to be directly tested; these were estimated using maximum likelihood with bootstrap confidence intervals (2,000 resamples) [Cumming and Finch, 2005]. Missing cognitive data were handled using full information maximum likelihood (FIML) in both approaches. Sex differences in path coefficients were evaluated by comparing bootstrap confidence intervals across groups.

#### 2.4.4 Sensitivity Analyses

Three sensitivity analyses assessed the robustness of sex difference findings to analytical choices.

##### 2.4.4.1 Site effects

Scanner site (UK Biobank assessment centre) can introduce systematic variance in volumetric measurements due to differences in scanner hardware, software versions, or acquisition protocols. Site effects were quantified using the intraclass correlation coefficient:

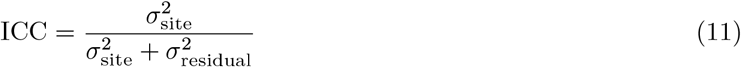

Site-adjusted sex difference effect sizes were computed using linear mixed-effects models with a fixed effect for sex and a random intercept for scanner site:

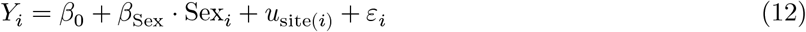

where 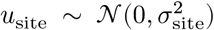 and 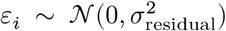.Cohen’s d was approximated as 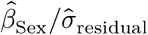 with confidence intervals derived from the Wald approximation.

##### 2.4.4.2 Hemisphere comparison

Left and right hippocampus and lateral ventricle volumes (residualized against ICV) were analyzed separately to assess whether sex differences are lateralized. Cohen’s d was computed for each hemisphere independently, with effect size comparisons based on overlapping 95% confidence intervals. Non-overlapping intervals indicate a statistically significant difference in effect magnitude between hemispheres.

##### 2.4.4.3 Inclusion of psychiatric diagnoses

To assess robustness, sex difference analyses were repeated in an expanded sensitivity sample that retained participants with F-code diagnoses (F00-F99; N = 30,032) while still excluding G and Q0 codes. The impact of psychiatric exclusion was quantified as Δ*d* = *d*_primary_ - *d*_sensitivity_.

## 3 Results

### 3.1 Sample Characteristics

The primary sample comprised 27,680 participants (15,555 female, 12,125 male; 56.2% female) aged 44.6–82.8 years (Table 1). Females were slightly younger on average (M = 62.9, SD = 7.3) than males (M = 64.1, SD = 7.7). Males had substantially larger ICV than females, consistent with the well-documented sex difference in head size. The England subsample (N = 25,172), ICV-matched subsample (N = 9,500; 4,750 pairs), and longitudinal subsample (N = 2,935) showed comparable age and ICV distributions to the primary sample within their respective selection constraints.

### 3.2 Sex Differences Across Adjustment Methods

#### 3.2.1 HVR Self-Normalizing Properties

Unadjusted HC showed strong positive correlation with ICV (r = 0.605), while proportions-adjusted HC showed moderate negative correlation (r = -0.39), indicating over-correction (Table 2). Residualization eliminated ICV association by construction (r ≈ 0). HVR’s correlation with ICV (r = -0.252) was smaller in magnitude than raw HC (r = 0.605) but remained negative, unlike residualized HC (r ≈ 0). This represents a 58% reduction in absolute ICV association. The negative direction mirrors the LV-ICV relationship (r = 0.462), indicating that the HVR-ICV association reflects the ventricular component’s scaling with head size.

**Table 2:** Brain measure correlations with ICV. An ideal self-normalizing measure shows r(ICV) near zero.

| Measure | $r(\text{ICV})$ | 95% CI |
| --- | --- | --- |
| Hippocampus |  |  |
| Unadjusted | 0.605 | [0.597, 0.612] |
| Proportions | -0.390 | [-0.400, -0.380] |
| Stereotaxic | -0.323 | [-0.333, -0.312] |
| Residualized | 0.000 | [-0.012, 0.012] |
| Lateral Ventricles |  |  |
| Unadjusted | 0.462 | [0.452, 0.471] |
| Proportions | 0.128 | [0.116, 0.139] |
| Stereotaxic | 0.134 | [0.122, 0.146] |
| Residualized | 0.000 | [-0.012, 0.012] |
| Hippocampal-to-Ventricle Ratio |  |  |
| Self-normalizing | -0.252 | [-0.263, -0.241] |

#### 3.2.2 Sex Differences in Brain Measures

Sex differences varied across brain measures and adjustment methods (N = 27,680; Figure 2).

**Figure 2:**
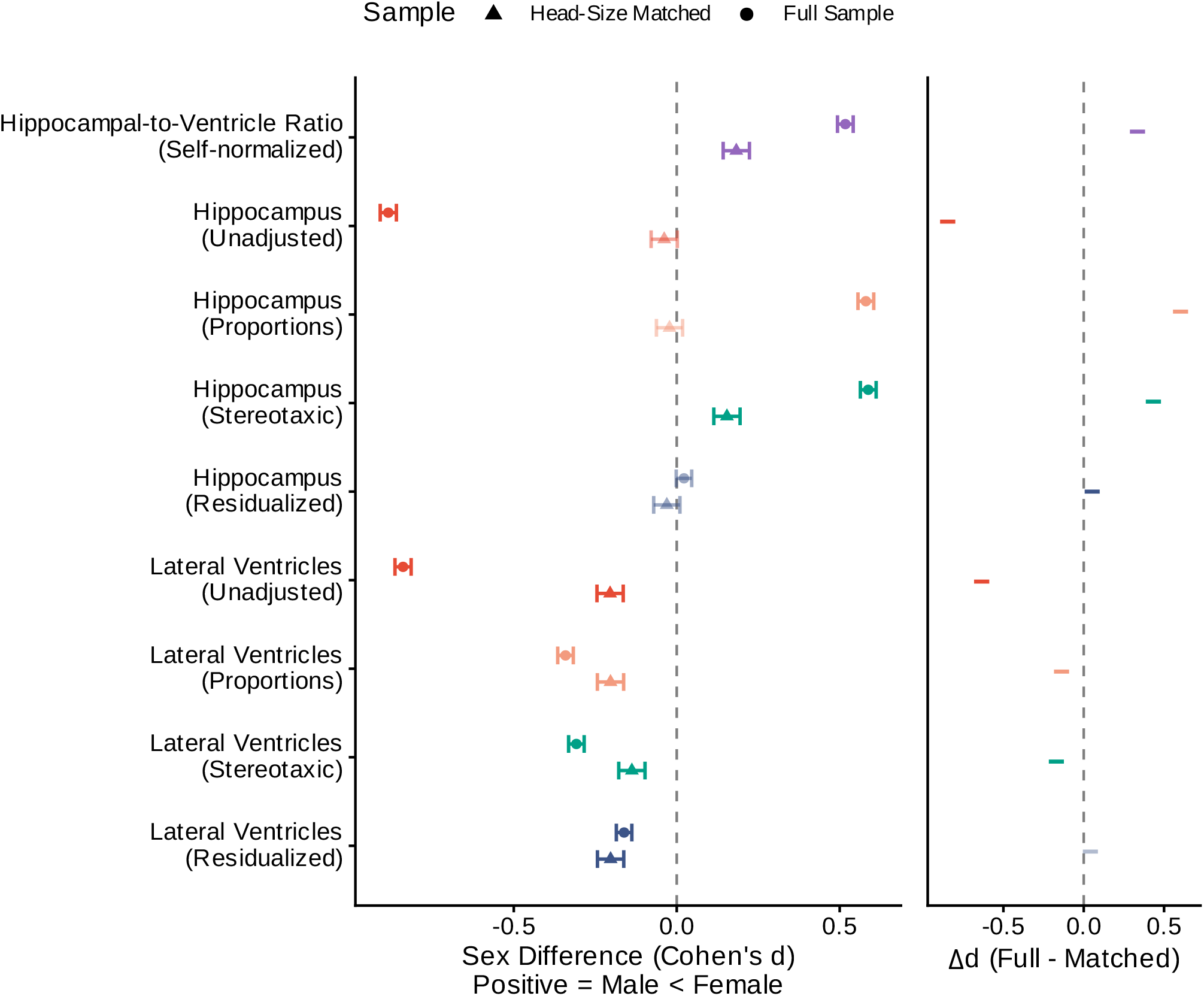
Sex differences by adjustment method. Circles = full sample; triangles = ICV-matched subsample. Faded points indicate non-significant effects (95% confidence interval crosses zero). Right panel shows Δ*d* = *d*_full_ − *d*_matched_, representing the change in effect size after removing head-size confounding; values near zero indicate head-size-independent effects.

##### 3.2.2.1 Hippocampal Volume

HC sex differences reversed direction depending on the adjustment method. Unadjusted volumes showed d = -0.89 [95% CI: -0.91, -0.86], indicating males had larger absolute volumes. Proportions adjustment reversed this to d = 0.58 [0.56, 0.6] with females larger. Stereotaxic adjustment yielded d = 0.59 [0.56, 0.61], and residualized volumes showed a negligible effect (d = 0.02 [0, 0.05]). The effect size range across methods spanned 1.47 standard deviations.

##### 3.2.2.2 Lateral Ventricular Volume

Unlike HC, lateral ventricular sex differences were consistently negative (males > females) across all adjustment methods, though effect magnitudes were attenuated by head-size correction. Unadjusted LV showed d = -0.84 [-0.86, -0.82], similar in magnitude to unadjusted HC. After adjustment, all methods retained the male advantage: proportions d = -0.34 [-0.37, -0.32], stereotaxic d = -0.31 [-0.33, -0.28], and residualized d = -0.16 [-0.19, -0.14]. The directional consistency across methods contrasts with the reversals observed for HC.

##### 3.2.2.3 HVR

HVR showed a consistent female advantage: d = 0.52 [95% CI: 0.49, 0.54]. The combination of males having relatively smaller hippocampi (after head-size adjustment) and larger ventricles produces this female advantage in HVR. Distribution overlap between sexes across adjustment methods is shown in Supporting Information (Figure S2).

##### 3.2.2.4 Matched-Pair Comparison

Matching on age and ICV (N = 9,500; 4,750 pairs) revealed which sex differences persist after removing head-size confounding (Table 3). HC effects collapsed to near-zero (unadjusted d = -0.04), confirming that HC sex differences are driven by head-size variation. LV effects were attenuated but remained significant (unadjusted d = -0.2), indicating that males have larger ventricles even at equivalent head size. HVR was reduced but persisted (d = 0.18 [0.14, 0.22]), consistent with the female HVR advantage being independent of head-size confounding.

**Table 3:** Full vs. ICV-matched subsample effect sizes. Matching on age (±1 yr) and ICV (±25 cc) eliminates head-size confounding.

| Adjustment | Full Sample |  | Matched Sample |  | Difference |  |
| --- | --- | --- | --- | --- | --- | --- |
| | d | 95% CI | d | 95% CI | $\Delta d$ | 95% CI |
| Hippocampus |  |  |  |  |  |  |
| Unadjusted | -0.885 | [-0.910, -0.860] | -0.038 | [-0.079, 0.002] | -0.847 | [-0.894, -0.799] |
| Proportions | 0.580 | [0.556, 0.605] | -0.022 | [-0.062, 0.018] | 0.602 | [0.556, 0.649] |
| Residuals | 0.022 | [-0.002, 0.046] | -0.030 | [-0.071, 0.010] | 0.053 | [0.006, 0.099] |
| Stereotaxic | 0.588 | [0.563, 0.612] | 0.154 | [0.114, 0.194] | 0.434 | [0.387, 0.481] |
| Lateral Ventricles |  |  |  |  |  |  |
| Unadjusted | -0.840 | [-0.865, -0.815] | -0.204 | [-0.245, -0.164] | -0.635 | [-0.683, -0.588] |
| Proportions | -0.341 | [-0.365, -0.317] | -0.203 | [-0.244, -0.163] | -0.138 | [-0.185, -0.091] |
| Residuals | -0.162 | [-0.185, -0.138] | -0.203 | [-0.243, -0.163] | 0.041 | [-0.005, 0.088] |
| Stereotaxic | -0.308 | [-0.332, -0.284] | -0.138 | [-0.178, -0.097] | -0.170 | [-0.217, -0.123] |
| Hippocampus-to-Ventricle ratio |  |  |  |  |  |  |
| Self-normalizing | 0.517 | [0.493, 0.542] | 0.183 | [0.142, 0.223] | 0.335 | [0.288, 0.382] |

### 3.3 Age-Related Patterns and Normative Modeling

#### 3.3.1 Age × Sex Interactions

Cross-sectional age patterns (N = 27,680) differed by sex (Figure 3). For HVR, the age × sex interaction was significant (*β* = -0.001, SE = 0, t = -16.58, p = 2e-61), with males showing 1.66× steeper cross-sectional age-related differences. The female HVR advantage *widened* with age—effect sizes increased from d = 0.23 at ages 45-55 to d = 0.66 at ages 75+.

**Figure 3:**
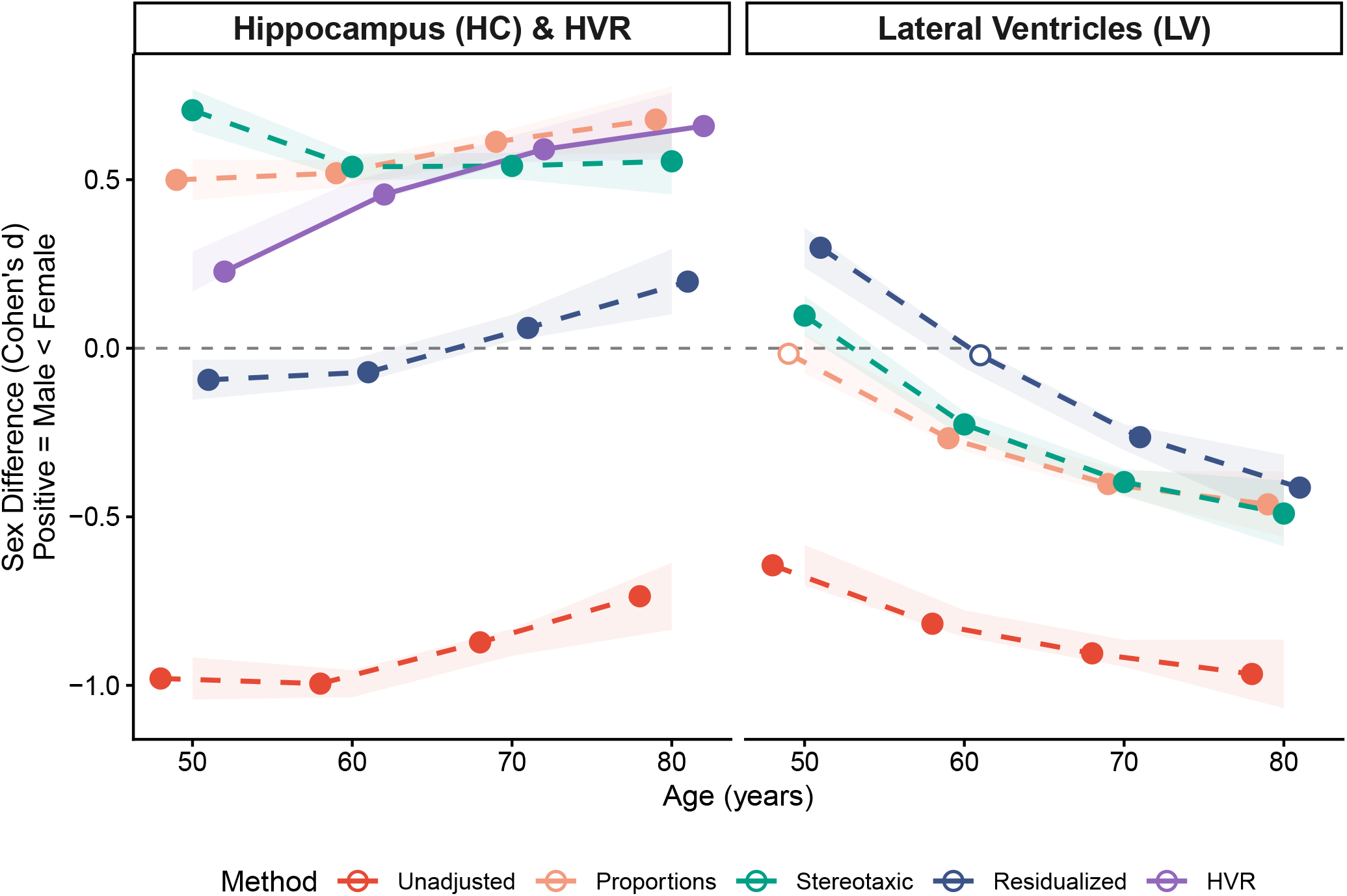
Age-related patterns in sex differences. Solid line = hippocampal-to-ventricle ratio; dashed lines = hippocampal/ventricular volume. White-filled points = non-significant (95% confidence interval crosses zero).

Age × sex interactions were significant for all three measures. HC showed males with 1.63× steeper cross-sectional differences (p = 3.1e-10). LV showed the largest sex disparity, with males exhibiting 1.87× steeper ventricular expansion (p = 4.6e-80). The greater sex disparity in ventricular age patterns compared to hippocampal age patterns explains the intermediate HVR ratio (1.66×), as HVR captures both processes.

#### 3.3.2 Normative Centile Curves

Figure 4 presents sex-specific normative centiles for HVR, hippocampal volume, and lateral ventricular volume. Across the age range, females showed higher HVR values at all percentile levels; the median female HVR at age 55 corresponded approximately to the 75th percentile for males of the same age. HC and LV centile curves are generated from GAMLSS models that include ICV as a covariate, providing head-size adjusted norms without pre-residualizing the data. Males show greater ventricular variability at older ages, potentially reflecting greater heterogeneity in aging patterns.

**Figure 4:**
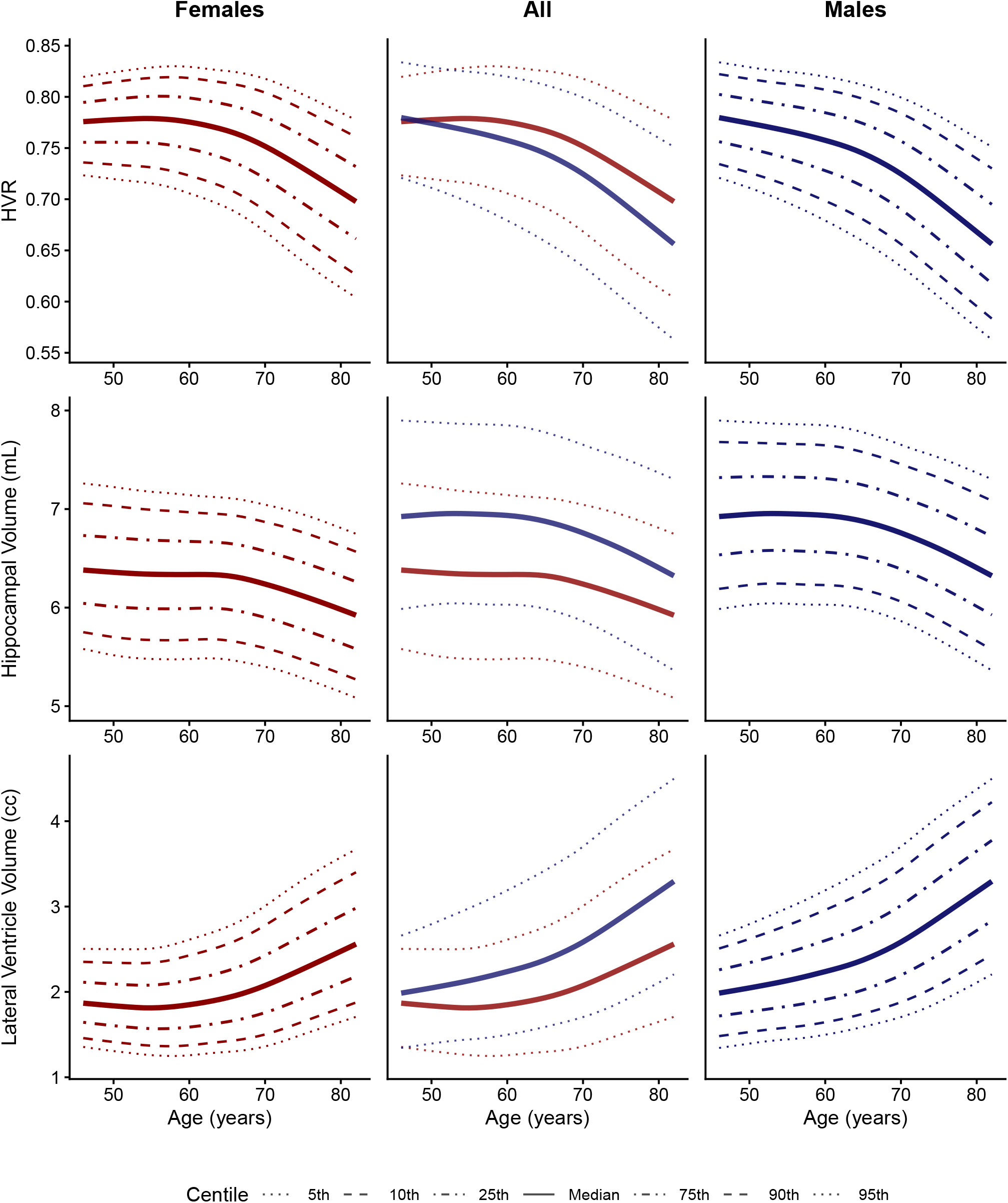
Sex-specific normative centile curves. Hippocampal-to-ventricle ratio (top), hippocampal volume (middle), and lateral ventricular volume (bottom). Left and right columns show female and male centiles separately; the centre column overlays both sexes for direct comparison. Hippocampal and ventricular volume models include intracranial volume as a covariate.

Calibration plots (Supporting Information, Figure S1) assess whether the normative models are well-specified: for each nominal centile (e.g., the 10th percentile curve), the plot shows what proportion of observations actually fall below that curve. A well-calibrated model produces points along the diagonal (observed proportion = expected proportion). Our models closely track the diagonal across all centiles, indicating that the fitted distributions accurately capture the true variability in the data.

##### 3.3.2.1 Temporal Stability (Longitudinal Validation)

All brain measures demonstrated high test-retest reliability across the longitudinal follow-up (Table S3), with Pearson correlations > 0.95 and ICCs > 0.93 for all measures. HVR showed slightly lower reliability (r = 0.952, ICC = 0.933) compared to HC (r = 0.964, ICC = 0.959) and LV (r = 0.958, ICC = 0.946), consistent with the ratio measure amplifying small measurement errors in either component. These reliabilities support the use of these normative models for individual-level assessment.

##### 3.3.2.2 Clinical Normative Centile Tables

Sex-specific normative centile tables for HVR, hippocampal volume, and lateral ventricular volume are provided in Supporting Information (Tables S4–S9) to enable clinical interpretation of individual measurements at 5-year age intervals. These tables allow clinicians to determine an individual’s percentile rank given their age, sex, and brain measure.

GAMLSS model coefficients for the transfer models (excluding site random effects) are provided in Table S10 and Table S11. These transfer models enable researchers to compute individual centile scores in external datasets where UK Biobank site identifiers are not applicable.

### 3.4 Brain-Cognition Associations

SEM analyses were restricted to the England subsample (N = 25,172) to ensure comparability of the Index of Multiple Deprivation (IMDP) covariate.

#### 3.4.1 Measurement Invariance

The cognitive bifactor model showed acceptable fit at all invariance levels, with ΔCFI ≤ 0.01 and ΔRMSEA ≤ 0.015 at each step (Supporting Information, Table S12). Strict measurement invariance was supported, validating cross-sex comparisons of factor scores and structural paths.

#### 3.4.2 Pooled Model Results

General cognition (*g*) was positively associated with all three brain measures (Table 4): unadjusted HC (*β* = 0.04 [95% CI: 0.023, 0.056]), HVR (*β* = 0.043 [0.027, 0.058]), and HC_RES_ (*β* = 0.028 [0.014, 0.042]), all p < .001. Confidence intervals overlapped substantially, indicating no significant differences in predictive validity between measures. HVR showed numerically the strongest association with *g*, despite requiring no head-size adjustment.

**Table 4:** Pooled brain-cognition path coefficients. Beta = standardized path coefficient; CI from 2000 bootstrap resamples. Memory-specific factor paths not shown due to factor collapse.

| Path | $\beta$ | 95% CI | p |
| --- | --- | --- | --- |
| HVR |  |  |  |
| →g | **0.043** | **[0.027, 0.058]** | **5.73e-08** |
| →Speed<sub>s</sub> | 0.022 | [-0.000, 0.044] | 0.055 |
| HC (Residualized) |  |  |  |
| →g | **0.028** | **[0.014, 0.042]** | **1.30e-04** |
| →Speed<sub>s</sub> | **0.033** | **[0.012, 0.054]** | **0.002** |
| HC (Unadjusted) |  |  |  |
| →g | **0.040** | **[0.023, 0.056]** | **2.65e-06** |
| →Speed<sub>s</sub> | **0.041** | **[0.017, 0.066]** | **7.35e-04** |

For processing speed (Speed_s_), HC showed the strongest effect (*β* = 0.041, p < .001), followed by HC_RES_ (*β* = 0.033, p = 0.002), with HVR trending but not significant (*β* = 0.022, p = 0.055). The memory-specific factor (Memory_s_) could not be reliably estimated in either sex and is therefore not reported here (see Table S15 for factor loadings).

#### 3.4.3 Sex-Stratified Results

Figure 5 displays sex-stratified path coefficients from multi-group models (detailed in Table S13). The multigroup analyses revealed sex differences in brain-cognition associations.

**Figure 5:**
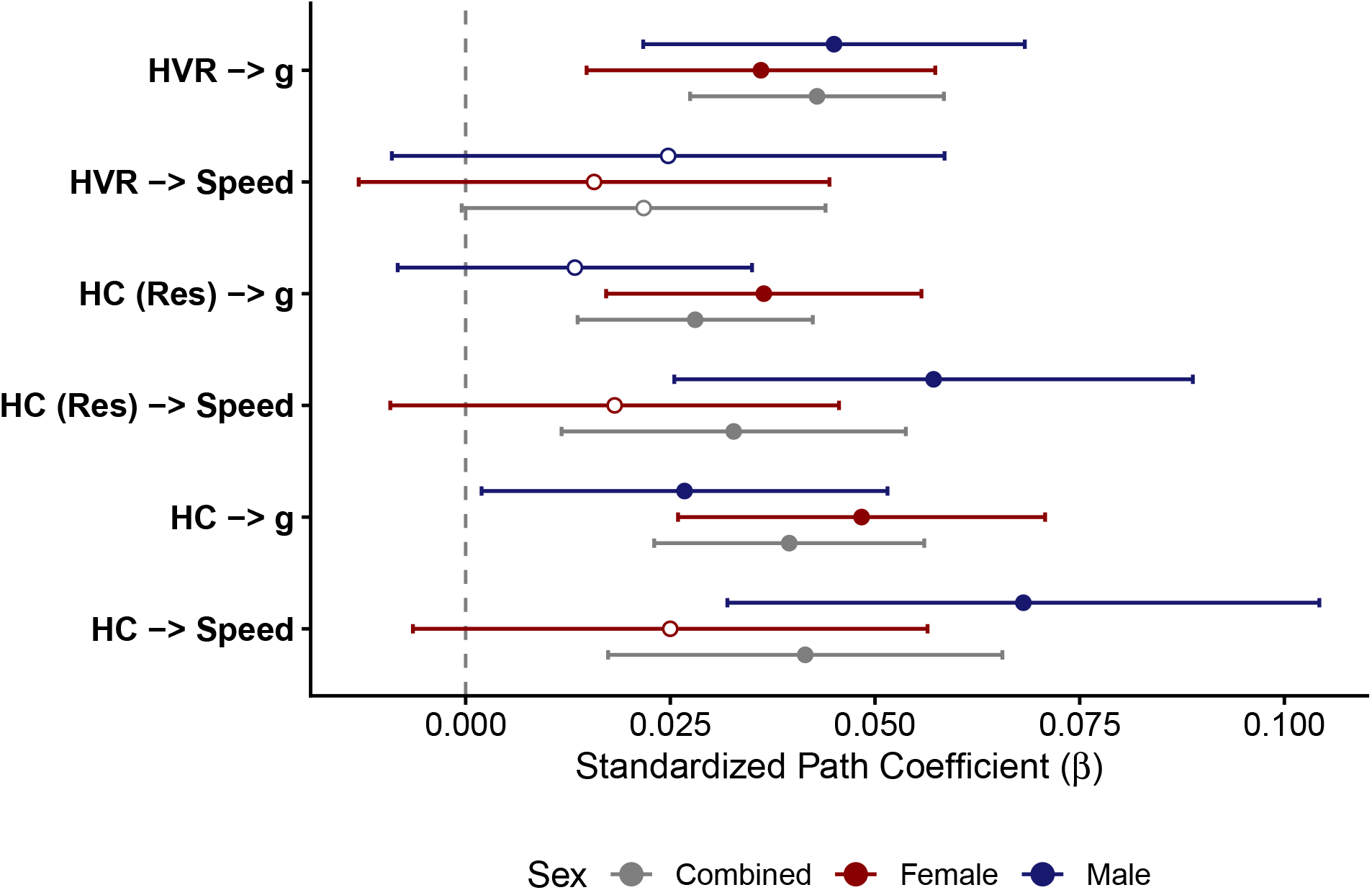
Brain-cognition paths by sex. Points represent standardized coefficients; lines show 95% bootstrap confidence intervals. White-filled points indicate non-significant effects.

For general cognition (*g*), HC_RES_ ⟶ *g* was not significant in males (p = 0.227) while remaining significant in females. In contrast, HVR showed the opposite pattern: males exhibited stronger HVR-cognition associations (*β* = 0.045) than females (*β* = 0.036), with both significant. For processing speed (Speed_s_), males showed significant HC and HC_RES_ associations (*β* = 0.068 and 0.057, respectively) while females showed none, suggesting sex-differential neural substrates for processing speed performance. Neither sex showed significant HVR ⟶ Speed_s_ associations.

All SEM models demonstrated good fit according to conventional criteria (Table S14; Hu and Bentler [1999]). Full model parameters including factor loadings and covariate effects are reported in Table S15 and Table S16.

##### 3.4.3.1 Non-Linear Age Effects

As described in Methods, quadratic age terms were included in HVR and HC_RES_ models but not in the HC model due to multicollinearity; fit indices for the HC model (Table S14) therefore reflect a linear-age specification. All quadratic age effects were significantly negative (all p < .001; Table S16), indicating that cross-sectional differences between older and younger participants were disproportionately larger at older ages. HVR showed a stronger quadratic effect (*β* = -0.134) than HC_RES_ (*β* = -0.075; Table S16), consistent with HVR capturing both hippocampal and ventricular age-related changes, both of which accelerate at older ages. Cognitive factors showed comparable quadratic effects across models (*g*: *β* ≈ -0.078; Speed_s_: *β* ≈ -0.046).

### 3.5 Sensitivity Analyses

#### 3.5.1 Site-Adjusted Effect Sizes

Site ICCs ranged from 0.004 to 0.012 (Supporting Information, Table S17), indicating that scanner site explained less than 1.5% of total variance in all brain measures. Effect sizes remained consistent after site adjustment.

#### 3.5.2 Hemisphere Comparison

Sex difference effect sizes were largely symmetrical across hemispheres for the individual volumes (Table S18, Figure S3). Residualized hippocampal volume showed significant effects bilaterally (left d = −0.14, right d = −0.16) with overlapping confidence intervals, indicating no lateralized sex difference. Lateral ventricles similarly showed a male advantage in both hemispheres with nearly identical effect sizes (left d = 0.48, right d = 0.48). However, HVR showed a lateralized pattern: the right hemisphere exhibited a significant female advantage (d = 0.05 [0.03, 0.07]), while the left hemisphere effect was non-significant (d = −0.01 [−0.03, 0.02]).

#### 3.5.3 Inclusion of Psychiatric Diagnoses (F-Codes)

Effect sizes remained stable when including participants with psychiatric diagnoses (N = 30,032; Table S19, Figure S4), demonstrating robustness of findings to exclusion criteria.

## 4 Discussion

### 4.1 Summary of Findings

In 27,680 UK Biobank participants, we evaluated HVR as a self-normalizing measure that bypasses conventional head-size correction. HVR showed weak correlation with ICV, substantially reducing head-size dependence compared to unadjusted HC. Hippocampal sex differences ranged from d = -0.89 (males larger) to d = 0.58 (females larger) depending solely on the adjustment method and age, whereas HVR consistently showed a female advantage replicated in the ICV-matched subsample. Males showed steeper age-related HVR differences (1.66× female slope), with the female advantage widening at older ages (d = 0.23 at 45-55y to d = 0.66 at 75+y). HVR predicted general cognition comparably to hippocampal volume.

### 4.2 Sex Differences Across Adjustment Methods

HVR achieves head-size independence through its mathematical construction: both numerator (HC) and denominator (HC + LV) scale with head size, so their ratio largely normalizes out this confound. Schoemaker et al. [2019] demonstrated that HVR yielded stronger correlations with both age and memory performance than absolute hippocampal volume, supporting its utility as an integrity index. In the present sample, HVR achieved a 58% reduction in ICV correlation compared to unadjusted HC (r = -0.252 vs. r = 0.605). This approach complements other ratio measures in clinical neuroimaging, such as the Evans’ index [Currà et al., 2019], and avoids the pitfalls of proportional ICV correction when brain structures scale non-isometrically with head size [Pintzka et al., 2015; Wang et al., 2024]. The residual negative HVR-ICV correlation likely reflects ventricular scaling: the hippocampus scales hypoallometrically with ICV while lateral ventricles scale near-isometrically [Jong et al., 2017], so larger heads have disproportionately larger ventricles relative to hippocampal volume, naturally producing lower HVR. Unlike residualization, which removes all ICV association by construction, HVR preserves morphological variation related to how ventricular space scales with brain size.

The reversal of HC sex differences across adjustment methods—from d = -0.89 (males larger, unadjusted) to d = 0.58 (females larger, proportions-adjusted), a swing of 1.47 SD—indicates these differences reflect head-size confounding rather than biology. This over-correction arises because the hippocampus scales hypoallometrically with ICV (*α* = 0.60; Jong et al. [2017]), so dividing by ICV systematically overestimates relative volume in smaller heads [Pintzka et al., 2015; Voevodskaya et al., 2014]. The pattern replicates across multiple cohorts [Fjell et al., 2009; Pintzka et al., 2015; Voevodskaya et al., 2014], and Ritchie et al. [2018] reported that unadjusted hippocampal sex differences in the UK Biobank dropped to near-zero after regression-based adjustment. The near-zero residualized effect (d = 0.022) suggests no meaningful sex difference after appropriate statistical control. While total hippocampal volume sex differences appear methodologically determined, sex-specific trajectories in hippocampal subregions [Mu et al., 2020] and structural connectivity patterns [Ingalhalikar et al., 2014] suggest meaningful differences may exist at finer spatial scales. Lateral ventricles, by contrast, showed a consistent male advantage across all methods (Figure 2). Because the LV-ICV regression has a negative y-intercept, proportional correction under-corrects rather than over-corrects ventricular volumes—the opposite of what occurs for HC [Pintzka et al., 2015]. The male advantage persists or increases after correction [Fjell et al., 2009], and in our large sample, residualized LV differences remained significant (d = -0.16), suggesting that null findings in smaller studies may reflect insufficient power. The consistent male ventricular advantage contrasts sharply with the method-dependent HC reversals, indicating this sex difference is not a head-size artifact.

The matched-pair analysis (Table 3) provides the strongest test of head-size independence. Matching on age and ICV eliminates head-size variation without the mathematical assumptions of statistical correction [Brzezinski-Rittner et al., 2025]. When matched, the HC sex difference collapsed to d = -0.038 (effectively zero), replicating prior matched-sample findings [Pintzka et al., 2015; Voevodskaya et al., 2014], while LV retained a male advantage (d = -0.2) and HVR retained a significant female advantage (d = 0.183). The HVR effect decreased 65% from full to matched samples, indicating that most of the sex difference is attributable to head-size variation, but a significant residual persists. Since HVR = HC/(HC+LV), the female advantage arises from two converging factors: comparable hippocampal volumes after size matching and relatively smaller female ventricles. This convergence—HC collapsing while LV and HVR effects survive ICV matching—confirms that the female HVR advantage reflects genuine sex differences in the hippocampus-to-ventricle balance, not head-size confounding.

### 4.3 Age-Related Patterns and Normative Modeling

Males showed steeper cross-sectional differences across all measures: 1.66× for HVR (p = 2e-61), 1.63× for HC, and 1.87× for LV, consistent with steeper male subcortical decline reported by Ritchie et al. [2018] and steeper male CSF expansion by Fjell et al. [2009]. The greater sex disparity in ventricular differences (1.87×) compared to hippocampal differences (1.63×) explains HVR’s intermediate ratio: HVR captures both processes, and steeper male ventricular expansion amplifies hippocampal differences. Males also show earlier ventricular abnormality signatures, with ventricular enlargement accelerating in the seventies [Currà et al., 2019], consistent with the steeper male HVR differences observed here. While the cross-sectional design precludes identifying a specific onset age, our age-stratified effect sizes show the male-female HVR divergence is already present in the youngest bin (ages 45–55, d = 0.23) and widens monotonically with age (d = 0.66 at ages 75+), suggesting the divergence begins before midlife. However, Brzezinski-Rittner et al. [2025] found that hippocampal age × sex interactions disappeared in a TIV-matched subsample, suggesting that some sex-differential aging trajectories may be byproducts of initial brain size differences.

The prognostic relevance of age-pattern differences is supported by evidence that steeper hippocampal volume loss predicts dementia conversion [Rajagopal et al., 2024]. Cumulative estrogen exposure may contribute to the female advantage, as longer reproductive spans are associated with younger brain ages [Luders et al., 2025] and estrogen promotes hippocampal neurogenesis and synaptic plasticity [Jett et al., 2022]. However, the female advantage is not universal: female APOE *ε*4 carriers show faster hippocampal loss than male carriers in aMCI [Park et al., 2025], indicating that genetic risk can modulate typical sex-differential patterns. Other large studies have found no significant sex differences in hippocampal age slopes [Fjell et al., 2009; Pintzka et al., 2015], underscoring that the presence of age × sex interactions depends on sample size, age range, and head-size adjustment method. These cross-sectional patterns cannot establish true individual trajectories and may be confounded by cohort effects or differential survival; longitudinal validation is essential.

The sex-specific centile curves (Figure 4) extend normative brain charts [Bethlehem et al., 2022; Dima et al., 2022; Ge et al., 2024] by providing the first sex-specific reference values for HVR. Our GAMLSS approach aligns with Bethlehem et al. [2022] and Dima et al. [2022], flexibly modeling both non-linear age trends and age-related variability changes. Sex-specific norms are essential to avoid misclassification, consistent with recommendations from Williams et al. [2021] and Ge et al. [2024], who found that sex accounts for a considerable proportion of morphological variance. For residualized HC, sex differences in centiles are minimal, consistent with the near-zero Cohen’s d for residualized volumes, supporting combined norms when proper ICV adjustment is applied. For LV, males show greater ventricular variability at older ages, potentially reflecting greater heterogeneity in aging-related patterns. Well-calibrated centile estimates (Figure S1) have demonstrated clinical utility in detecting case-control differences and predicting diagnostic transition from MCI to Alzheimer’s disease [Bethlehem et al., 2022; Dima et al., 2022]. HVR showed slightly lower test-retest reliability (ICC > 0.93; Table S3) than HC or LV alone over a mean 2.7-year follow-up. Because this interval spans real aging-related change, the lower ICC may partly reflect HVR’s sensitivity to concurrent hippocampal atrophy and ventricular expansion rather than measurement noise alone — a hypothesis that dedicated longitudinal studies with shorter retest intervals could directly test.

### 4.4 Brain-Cognition Associations

Strict measurement invariance for the cognitive bifactor model (ΔCFI ≤ 0.01 and ΔRMSEA ≤ 0.015 at each level; Table S12) confirmed that the same cognitive constructs (*g*, Memory_s_, Speed_s_) exist in both sexes with equivalent loadings, intercepts, and residual variances [Chen, 2007; Putnick and Bornstein, 2016], validating cross-sex comparison of brain-cognition paths. In pooled models, HVR predicted general cognition comparably to HC without requiring head-size adjustment (HVR ⟶ *g β* = 0.043 vs. HC ⟶ *g β* = 0.04, overlapping CIs), with all effects small—consistent with the modest brain-cognition associations typically observed in large healthy samples [Ritchie et al., 2018; Wang et al., 2024]. The restricted cognitive range in healthy volunteer samples likely attenuates these associations [Bachmann et al., 2023]. Multigroup analyses revealed sex-differential patterns: HC_RES_ predicted cognition in females but not males (*β* = 0.013, p = 0.227), consistent with stronger female hippocampal-cognition associations reported elsewhere [Jockwitz et al., 2024; Zheng et al., 2017]. Raw HC showed significant associations in both sexes but with a female advantage (*β* = 0.048 vs. 0.027). HVR showed the opposite pattern, with males exhibiting stronger associations (*β* = 0.045 vs. 0.036), though both sexes reached significance.

These sex-differential patterns indicate that HVR is not simply “residualized HC by another means.” Unlike HC_RES_, which lost predictive validity in males, HVR maintained significant associations in both sexes— paralleling Schoemaker et al. [2019] and Fernandez-Lozano et al. [2025], who found that HVR predicted cognition comparably or better than HC alone. The differential patterns between HC_RES_ and HVR across sexes suggest HVR captures distinct information related to hippocampus-ventricle coupling. HVR also accommodated quadratic age specifications that raw HC could not (the HC + ICV + age^2^ combination produced multicollinearity), allowing detection of accelerating age-related patterns (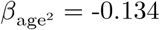 for HVR vs. -0.075 for HC_RES_). The stronger quadratic effect for HVR reflects its capture of both hippocampal and ventricular age-related differences. The ventricular component likely contributes cognitive-relevant variance beyond medial temporal integrity, as ventricular volume has been associated with executive function rather than episodic memory [Bachmann et al., 2023].

### 4.5 Sensitivity Analyses

Low site ICCs (all < 0.05; Table S17) confirmed minimal scanner site contribution to variance, and effect sizes were stable after site adjustment; the ratio construction of HVR did not amplify site-related noise. Sex difference effect sizes were symmetrical across hemispheres for HC and LV; HVR showed a small lateralized pattern (right d = 0.05 vs. left d = −0.01), though the small magnitude and potential contribution of hemispheric segmentation biases [Fernandez-Lozano et al., 2025] suggest bilateral HVR remains most robust for clinical application. Including participants with psychiatric diagnoses changed effect sizes minimally (Δd < 0.02; Table S19, Figure S4), consistent with the low prevalence of psychiatric diagnoses in this healthy volunteer cohort [Fry et al., 2017].

### 4.6 Limitations

Design and sample constraints include the cross-sectional design, which cannot distinguish age effects from cohort effects or establish individual change trajectories, and HVR’s conflation of hippocampal and ventricular contributions to observed differences. UK Biobank participants are healthier and more educated than the general population [Fry et al., 2017], limiting generalizability to clinical populations. Some cognitive tests had incomplete data, and complete-case analyses may introduce selection bias. APOE genotype was unavailable, precluding examination of gene × sex interactions relevant to dementia risk.

Methodologically, the study did not include power-corrected proportions allometric adjustment [Sanchis-Segura et al., 2020]. Imaging site clustering was not modeled in multigroup SEM because lavaan does not support this combination [Rosseel, 2012], though the minimal design effect across 4 sites makes sex comparisons conservative rather than biased. The bifactor model’s memory-specific factor could not be reliably estimated in either sex—a common occurrence in bifactor models [Burns et al., 2020; Eid et al., 2017]—indicating that memory variance was fully captured by *g*; brain-cognition paths involving Memory_s_ are therefore not reported. Ratios can exhibit non-normality and complex interpretation when numerator and denominator correlate differently with outcomes, though HVR showed approximately normal distributions here. All brain-cognition associations were small (*β* = 0.02–0.04), explaining less than 1% of variance in cognitive performance. While statistically significant given the large sample, the practical and clinical significance of these associations remains to be established.

### 4.7 Future Directions

The current findings motivate four lines of follow-up work. First, the UK Biobank repeat imaging subsample enables longitudinal validation of whether HVR predicts individual-level cognitive decline. Second, clinical validation in AD/MCI cohorts would assess HVR’s diagnostic utility compared to clinical populations studied previously [Fernandez-Lozano et al., 2025]. Third, longitudinal data could separate hippocampal and ventricular contributions to HVR changes, which the cross-sectional ratio conflates. Fourth, examining whether HVR × sex × APOE genotype predicts dementia conversion would address the gene × sex interactions that this study could not explore.

### 4.8 Conclusions

In 27,680 UK Biobank participants (56.2% female), hippocampal sex differences reversed direction across adjustment methods (d range: 1.5 SD), confirming their methodological determination. HVR demonstrated substantial head-size independence (r_ICV_ = -0.25; 58% reduction vs. raw HC) and a consistent female advantage replicated in the ICV-matched subsample (d = 0.18). Males showed 1.66× steeper age-related HVR differences (p = 2e-61); sex-specific normative centile curves are provided for research application, pending validation in clinical populations. HVR predicted general cognition comparably to hippocampal volume (HVR ⟶ *g*: *β* = 0.043 vs. HC ⟶ *g*: *β* = 0.04, overlapping CIs), maintained predictive validity in both sexes while residualized HC lost significance in males (p = 0.227), and supported more complex SEM specifications that raw HC could not.

## Author Contributions

S.F-L. conceived the study, developed the analysis pipeline, performed all analyses, and wrote the manuscript.

D.L.C. supervised the project, provided critical review, and approved the final manuscript.

## Acknowledgments

This research has been conducted using the UK Biobank Resource under Application Number 45551. This investigation was supported in part by an award from the International Progressive MS Alliance (award reference number PA-1412-02420), a grant from the Canadian Institutes of Health Research, and a donation from the Famille Louise & Andre Charron.

## Data Availability Statement

This research was conducted using UK Biobank data (Application 45551). Data are available to approved researchers through the UK Biobank Access Management System (https://www.ukbiobank.ac.uk/). Analysis code is available at https://github.com/soffiafdz/hvr-sex-differences.

## Ethics Approval Statement

UK Biobank received ethical approval from the North West Multi-centre Research Ethics Committee (REC reference: 16/NW/0274). All participants provided written informed consent.

