## Supporting Information for "Hippocampal-to-Ventricle Ratio as a Head-Size-Independent Biomarker: Sex Differences and Cognitive Associations in 27,680 UK Biobank Participants"

Sofia Fernandez-Lozano D. Louis Collins

#### Appendix A: Cognitive Battery and Covariates

Table S1: **Cognitive test battery.** UK Biobank imaging visit assessments with field identifiers.

| Test | Field ID | Description |
|---|---|---|
| Memory | | |
| Pairs Matching (round 1) | 399 | Number of errors in visual memory task |
| Pairs Matching (round 2) | 400 | Number of errors in delayed recall |
| Prospective Memory | 20018 | Correct execution of delayed instruction |
| Processing Speed | | |
| Reaction Time | 20023 | Mean response time (ms) |
| Symbol Digit Substitution | 20159 | Number correct in 60 seconds |
| Trail Making A | 6348 | Time to complete (s) |
| Executive Function | | |
| Trail Making B | 6350 | Time to complete (s) |
| Reasoning | | |
| Fluid Intelligence | 20016 | Number correct in 13 verbal-numerical items |
| Matrix Reasoning | 6373 | Number correct in non-verbal reasoning |
| Tower Rearranging | 21004 | Planning task score |
| Verbal | | |
| Vocabulary | 6364 | Number of synonyms identified |
| Numeric | | |
| Numeric Memory | 4282 | Maximum digit span |

Table S2: **SEM model covariates.** UK Biobank field identifiers. Age$^2$ included in HVR and HC-RES models only. IMDP available for England subsample only. Site modeled as random effect in GAMLSS; cluster-robust SE in pooled SEM.

| | Description | UK Biobank Field |
|---|---|---|
| Age | Age at imaging visit (years) | 21003 |
| Age$^2$ | Quadratic age term | Derived |
| ICV | Intracranial volume (cc) | Derived |
| IMDP | Index of Multiple Deprivation | 26410 |
| Education | Educational attainment (years) | 6138 |
| Site | UK Biobank assessment centre | 54 |

### Appendix B: Normative Centile Tables

Table S3: **Test-retest reliability of brain measures.** Mean follow-up interval of 2.7 years (SD = 1.1). N = 2,935 participants with both ses-2 and ses-3 imaging. ICC = intraclass correlation coefficient (two-way, absolute agreement); not to be confused with ICV (intracranial volume).

| Laterality | N | Pearson r | ICC |
|---|---|---|---|
| Hippocampal-to-Ventricle Ratio | | | |
| Bilateral | 2935 | 0.952 | 0.933 |
| Left | 2935 | 0.955 | 0.940 |
| Right | 2935 | 0.945 | 0.929 |
| Hippocampus | | | |
| Bilateral | 2935 | 0.964 | 0.959 |
| Left | 2935 | 0.961 | 0.956 |
| Right | 2935 | 0.961 | 0.957 |
| Lateral Ventricles | | | |
| Bilateral | 2935 | 0.958 | 0.946 |
| Left | 2935 | 0.959 | 0.948 |
| Right | 2935 | 0.953 | 0.943 |

Table S4: **HVR normative centiles for females.** Values by hemisphere at 5-year age intervals for clinical reference.

| | Percentile | | | | |
|---|---|---|---|---|---|
| Age | 10th | 25th | 50th | 75th | 90th |
| Bilateral | | | | | |
| 50 | 0.734 | 0.756 | 0.778 | 0.798 | 0.815 |
| 55 | 0.731 | 0.755 | 0.779 | 0.801 | 0.819 |
| 60 | 0.723 | 0.749 | 0.775 | 0.799 | 0.818 |
| 65 | 0.710 | 0.739 | 0.767 | 0.792 | 0.813 |
| 70 | 0.689 | 0.720 | 0.752 | 0.780 | 0.804 |
| 75 | 0.662 | 0.696 | 0.730 | 0.762 | 0.788 |
| 80 | 0.636 | 0.671 | 0.707 | 0.740 | 0.769 |
| Left | | | | | |
| 50 | 0.722 | 0.750 | 0.776 | 0.799 | 0.818 |
| 55 | 0.718 | 0.748 | 0.776 | 0.801 | 0.821 |
| 60 | 0.709 | 0.741 | 0.772 | 0.799 | 0.820 |
| 65 | 0.695 | 0.729 | 0.762 | 0.791 | 0.814 |
| 70 | 0.673 | 0.709 | 0.745 | 0.778 | 0.804 |
| 75 | 0.644 | 0.683 | 0.722 | 0.758 | 0.787 |
| 80 | 0.614 | 0.655 | 0.696 | 0.734 | 0.766 |
| Right | | | | | |
| 50 | 0.735 | 0.757 | 0.781 | 0.803 | 0.821 |
| 55 | 0.733 | 0.758 | 0.783 | 0.806 | 0.827 |
| 60 | 0.727 | 0.753 | 0.780 | 0.805 | 0.826 |
| 65 | 0.716 | 0.744 | 0.773 | 0.800 | 0.822 |
| 70 | 0.696 | 0.727 | 0.759 | 0.789 | 0.813 |
| 75 | 0.670 | 0.705 | 0.739 | 0.771 | 0.797 |
| 80 | 0.647 | 0.683 | 0.719 | 0.753 | 0.781 |

Table S5: **HVR normative centiles for males.** Values by hemisphere at 5-year age intervals for clinical reference.

| | Percentile | | | | |
|---|---|---|---|---|---|
| Age | 10th | 25th | 50th | 75th | 90th |
| Bilateral | | | | | |
| 50 | 0.726 | 0.749 | 0.774 | 0.797 | 0.817 |
| 55 | 0.714 | 0.740 | 0.767 | 0.792 | 0.813 |
| 60 | 0.698 | 0.728 | 0.758 | 0.785 | 0.807 |
| 65 | 0.680 | 0.713 | 0.745 | 0.774 | 0.798 |
| 70 | 0.656 | 0.690 | 0.725 | 0.757 | 0.784 |
| 75 | 0.626 | 0.661 | 0.698 | 0.733 | 0.764 |
| 80 | 0.595 | 0.630 | 0.668 | 0.706 | 0.740 |
| Left | | | | | |
| 50 | 0.714 | 0.744 | 0.773 | 0.800 | 0.821 |
| 55 | 0.699 | 0.731 | 0.762 | 0.791 | 0.815 |
| 60 | 0.682 | 0.717 | 0.751 | 0.783 | 0.809 |
| 65 | 0.663 | 0.701 | 0.738 | 0.772 | 0.800 |
| 70 | 0.638 | 0.678 | 0.718 | 0.754 | 0.785 |
| 75 | 0.608 | 0.648 | 0.690 | 0.729 | 0.763 |
| 80 | 0.577 | 0.617 | 0.660 | 0.701 | 0.738 |
| Right | | | | | |
| 50 | 0.727 | 0.751 | 0.777 | 0.803 | 0.825 |
| 55 | 0.717 | 0.744 | 0.772 | 0.799 | 0.821 |
| 60 | 0.705 | 0.735 | 0.765 | 0.793 | 0.816 |
| 65 | 0.688 | 0.720 | 0.753 | 0.782 | 0.807 |
| 70 | 0.665 | 0.699 | 0.733 | 0.766 | 0.793 |
| 75 | 0.635 | 0.670 | 0.708 | 0.743 | 0.773 |
| 80 | 0.603 | 0.640 | 0.680 | 0.717 | 0.750 |

Table S6: **HC normative centiles for females.** Values by hemisphere (ICV-controlled). Given significant lateralization (right HC > left HC in females), hemisphere-specific norms are provided.

| | Percentile | | | | |
|---|---|---|---|---|---|
| Age | 10th | 25th | 50th | 75th | 90th |
| Bilateral | | | | | |
| 50 | 5.701 | 6.011 | 6.359 | 6.711 | 7.031 |
| 55 | 5.671 | 5.989 | 6.339 | 6.684 | 6.992 |
| 60 | 5.673 | 5.991 | 6.336 | 6.672 | 6.968 |
| 65 | 5.668 | 5.983 | 6.324 | 6.654 | 6.943 |
| 70 | 5.589 | 5.900 | 6.239 | 6.573 | 6.869 |
| 75 | 5.469 | 5.777 | 6.118 | 6.458 | 6.764 |
| 80 | 5.330 | 5.639 | 5.981 | 6.321 | 6.626 |
| Left | | | | | |
| 50 | 2.788 | 2.962 | 3.153 | 3.342 | 3.511 |
| 55 | 2.768 | 2.949 | 3.142 | 3.329 | 3.492 |
| 60 | 2.765 | 2.946 | 3.138 | 3.323 | 3.482 |
| 65 | 2.767 | 2.944 | 3.132 | 3.312 | 3.468 |
| 70 | 2.729 | 2.903 | 3.090 | 3.271 | 3.429 |
| 75 | 2.669 | 2.841 | 3.029 | 3.212 | 3.375 |
| 80 | 2.594 | 2.769 | 2.958 | 3.143 | 3.306 |
| Right | | | | | |
| 50 | 2.868 | 3.031 | 3.211 | 3.393 | 3.557 |
| 55 | 2.855 | 3.021 | 3.203 | 3.382 | 3.540 |
| 60 | 2.861 | 3.026 | 3.204 | 3.375 | 3.525 |
| 65 | 2.856 | 3.020 | 3.196 | 3.366 | 3.513 |
| 70 | 2.814 | 2.978 | 3.155 | 3.327 | 3.478 |
| 75 | 2.756 | 2.918 | 3.096 | 3.270 | 3.424 |
| 80 | 2.692 | 2.854 | 3.030 | 3.204 | 3.357 |

Table S7: **HC normative centiles for males.** Values by hemisphere (ICV-controlled).

| | Percentile | | | | |
|---|---|---|---|---|---|
| Age | 10th | 25th | 50th | 75th | 90th |
| Bilateral | | | | | |
| 50 | 6.230 | 6.568 | 6.947 | 7.328 | 7.674 |
| 55 | 6.242 | 6.579 | 6.952 | 7.326 | 7.661 |
| 60 | 6.229 | 6.563 | 6.935 | 7.309 | 7.647 |
| 65 | 6.183 | 6.510 | 6.876 | 7.244 | 7.578 |
| 70 | 6.061 | 6.391 | 6.758 | 7.125 | 7.455 |
| 75 | 5.877 | 6.218 | 6.598 | 6.977 | 7.319 |
| 80 | 5.662 | 6.013 | 6.404 | 6.798 | 7.155 |
| Left | | | | | |
| 50 | 3.063 | 3.249 | 3.454 | 3.657 | 3.838 |
| 55 | 3.060 | 3.248 | 3.455 | 3.660 | 3.843 |
| 60 | 3.053 | 3.238 | 3.442 | 3.647 | 3.830 |
| 65 | 3.033 | 3.213 | 3.413 | 3.613 | 3.793 |
| 70 | 2.973 | 3.154 | 3.353 | 3.549 | 3.724 |
| 75 | 2.876 | 3.066 | 3.271 | 3.473 | 3.651 |
| 80 | 2.768 | 2.964 | 3.176 | 3.382 | 3.563 |
| Right | | | | | |
| 50 | 3.112 | 3.298 | 3.502 | 3.702 | 3.881 |
| 55 | 3.122 | 3.307 | 3.506 | 3.698 | 3.867 |
| 60 | 3.118 | 3.302 | 3.501 | 3.695 | 3.865 |
| 65 | 3.097 | 3.275 | 3.470 | 3.661 | 3.830 |
| 70 | 3.036 | 3.216 | 3.412 | 3.604 | 3.774 |
| 75 | 2.951 | 3.133 | 3.334 | 3.533 | 3.710 |
| 80 | 2.847 | 3.030 | 3.236 | 3.442 | 3.629 |

Table S8: **LV normative centiles for females.** Values by hemisphere (ICV-controlled).

| | Percentile | | | | |
|---|---|---|---|---|---|
| Age | 10th | 25th | 50th | 75th | 90th |
| Bilateral | | | | | |
| 50 | 1.413 | 1.604 | 1.837 | 2.092 | 2.342 |
| 55 | 1.370 | 1.569 | 1.813 | 2.080 | 2.341 |
| 60 | 1.375 | 1.588 | 1.850 | 2.141 | 2.428 |
| 65 | 1.422 | 1.650 | 1.932 | 2.247 | 2.560 |
| 70 | 1.501 | 1.757 | 2.074 | 2.428 | 2.780 |
| 75 | 1.651 | 1.924 | 2.269 | 2.660 | 3.055 |
| 80 | 1.811 | 2.107 | 2.476 | 2.891 | 3.307 |
| Left | | | | | |
| 50 | 0.690 | 0.792 | 0.920 | 1.067 | 1.217 |
| 55 | 0.672 | 0.779 | 0.912 | 1.064 | 1.218 |
| 60 | 0.675 | 0.791 | 0.936 | 1.102 | 1.270 |
| 65 | 0.703 | 0.826 | 0.984 | 1.164 | 1.349 |
| 70 | 0.746 | 0.885 | 1.062 | 1.263 | 1.467 |
| 75 | 0.823 | 0.975 | 1.168 | 1.389 | 1.614 |
| 80 | 0.914 | 1.079 | 1.287 | 1.523 | 1.763 |
| Right | | | | | |
| 50 | 0.680 | 0.784 | 0.910 | 1.045 | 1.176 |
| 55 | 0.653 | 0.762 | 0.894 | 1.037 | 1.176 |
| 60 | 0.656 | 0.769 | 0.908 | 1.060 | 1.209 |
| 65 | 0.676 | 0.796 | 0.942 | 1.104 | 1.261 |
| 70 | 0.714 | 0.845 | 1.007 | 1.186 | 1.363 |
| 75 | 0.786 | 0.923 | 1.096 | 1.291 | 1.488 |
| 80 | 0.851 | 1.000 | 1.185 | 1.391 | 1.596 |

Table S9: **LV normative centiles for males.** Values by hemisphere (ICV-controlled).

| | Percentile | | | | |
|---|---|---|---|---|---|
| Age | 10th | 25th | 50th | 75th | 90th |
| Bilateral | | | | | |
| 50 | 1.530 | 1.767 | 2.047 | 2.342 | 2.621 |
| 55 | 1.582 | 1.831 | 2.132 | 2.460 | 2.780 |
| 60 | 1.645 | 1.911 | 2.238 | 2.601 | 2.960 |
| 65 | 1.743 | 2.023 | 2.374 | 2.769 | 3.166 |
| 70 | 1.880 | 2.196 | 2.584 | 3.012 | 3.433 |
| 75 | 2.082 | 2.439 | 2.869 | 3.332 | 3.776 |
| 80 | 2.331 | 2.719 | 3.173 | 3.650 | 4.098 |
| Left | | | | | |
| 50 | 0.746 | 0.871 | 1.026 | 1.200 | 1.373 |
| 55 | 0.781 | 0.917 | 1.085 | 1.274 | 1.461 |
| 60 | 0.815 | 0.963 | 1.148 | 1.356 | 1.563 |
| 65 | 0.866 | 1.021 | 1.219 | 1.446 | 1.678 |
| 70 | 0.937 | 1.110 | 1.327 | 1.574 | 1.823 |
| 75 | 1.039 | 1.235 | 1.476 | 1.740 | 1.999 |
| 80 | 1.169 | 1.381 | 1.635 | 1.907 | 2.169 |
| Right | | | | | |
| 50 | 0.730 | 0.861 | 1.012 | 1.169 | 1.315 |
| 55 | 0.749 | 0.880 | 1.039 | 1.213 | 1.382 |
| 60 | 0.777 | 0.914 | 1.083 | 1.272 | 1.459 |
| 65 | 0.825 | 0.969 | 1.149 | 1.349 | 1.548 |
| 70 | 0.892 | 1.053 | 1.249 | 1.463 | 1.672 |
| 75 | 0.990 | 1.168 | 1.383 | 1.617 | 1.843 |
| 80 | 1.110 | 1.299 | 1.526 | 1.770 | 2.003 |

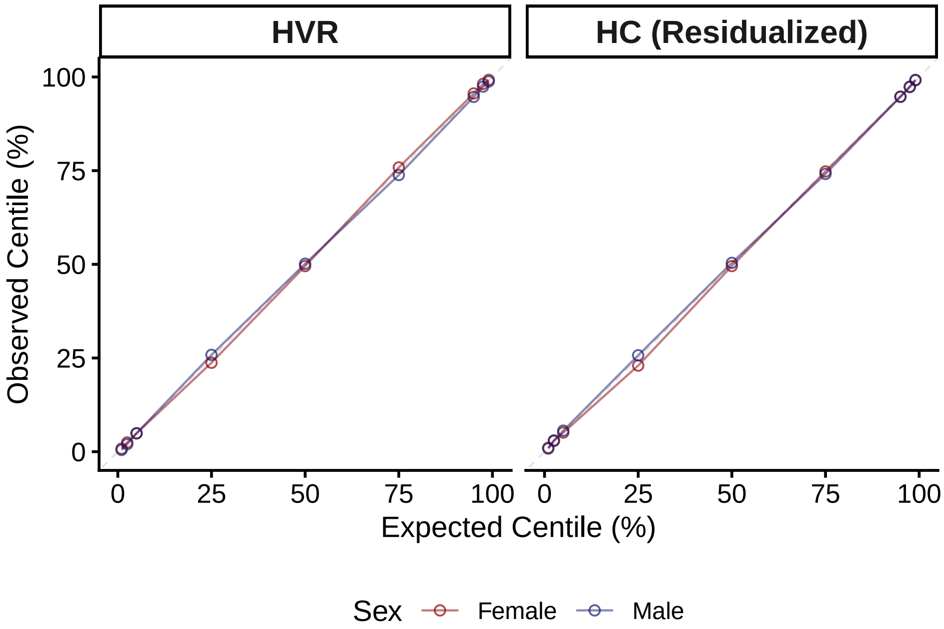


Figure S1: **GAMLSS normative model calibration.** Points represent observed vs. expected proportions in held-out test data. Diagonal line indicates perfect calibration.

#### Appendix C: Model Results

Table S10: **GAMLSS coefficients for HVR.** Transfer model without site. $\mu$ = location (mean), $\sigma$ = scale (variance), $\nu$ = skewness. Bold indicates p < .05.

| | Females | | | Males | | |
|---|---|---|---|---|---|---|
| Term | Estimate | SE | *p* | Estimate | SE | *p* |
| Location ($\mu$) | | | | | | |
| Age (cubic spline) | **-0.0020** | **0.0000** | **2.20e-16** | **-0.0032** | **0.0001** | **2.20e-16** |
| Education (years) | **-0.0007** | **0.0001** | **2.07e-08** | **-0.0008** | **0.0002** | **3.47e-07** |
| Intercept | **0.8994** | **0.0032** | **2.20e-16** | **0.9572** | **0.0040** | **2.20e-16** |
| Scale ($\sigma$) | | | | | | |
| Age (cubic spline) | **0.0203** | **0.0008** | **2.00e-16** | **0.0208** | **0.0008** | **2.00e-16** |
| Intercept | **-4.2618** | **0.0493** | **2.00e-16** | **-4.1385** | **0.0540** | **2.00e-16** |
| Shape ($\nu$) | | | | | | |
| Age (cubic spline) | **-0.0602** | **0.0160** | **1.70e-04** | **-0.0729** | **0.0149** | **9.91e-07** |
| Intercept | **7.6209** | **1.0487** | **3.85e-13** | **7.6470** | **0.9963** | **1.77e-14** |

Table S11: **GAMLSS coefficients for HC volume.** Transfer model without site. $\mu$ = location (mean), $\sigma$ = scale (variance), $\nu$ = skewness. Bold indicates p < .05.

| | Females | | | Males | | |
|---|---|---|---|---|---|---|
| Term | Estimate | SE | *p* | Estimate | SE | *p* |
| Location ($\mu$) | | | | | | |
| Age (cubic spline) | **-0.0092** | **0.0006** | **2.20e-16** | **-0.0161** | **0.0007** | **2.00e-16** |
| Education (years) | **0.0068** | **0.0015** | **8.26e-06** | **0.0048** | **0.0019** | **0.012** |
| Intercept | **2.6552** | **0.0698** | **2.20e-16** | **3.5113** | **0.0849** | **2.00e-16** |
| Intracranial Volume | **0.0027** | **0.0000** | **2.20e-16** | **0.0025** | **0.0000** | **2.00e-16** |
| Scale ($\sigma$) | | | | | | |
| Age (cubic spline) | -0.0002 | 0.0008 | 0.812 | **0.0023** | **0.0008** | **0.007** |
| Intercept | **-2.5101** | **0.0493** | **2.00e-16** | **-2.6603** | **0.0540** | **2.20e-16** |
| Shape ($\nu$) | | | | | | |
| Age (cubic spline) | 0.0068 | 0.0102 | 0.504 | 0.0012 | 0.0108 | 0.910 |
| Intercept | 0.8825 | 0.6465 | 0.172 | 0.8614 | 0.7023 | 0.220 |

Table S12: **Measurement invariance testing.** Cognitive bifactor model. Delta-CFI and Delta-RMSEA within recommended thresholds at each step supports strict invariance.

| Model | CFI | RMSEA | ΔCFI | ΔRMSEA |
|---|---|---|---|---|
| Configural | 0.963 | 0.059 | — | — |
| Metric (equal loadings) | 0.963 | 0.055 | -0.001 | -0.004 |
| Scalar (equal intercepts) | 0.957 | 0.057 | -0.006 | 0.002 |
| Strict (equal residuals) | 0.956 | 0.054 | -0.001 | -0.002 |

Table S13: **Brain-cognition paths by sex.** $\beta$ = standardized coefficient. Bold indicates p < .05. Multi-group SEM with sex-stratified parameter estimation.

| | Females | | | Males | | |
|---|---|---|---|---|---|---|
| Outcome | $\beta$ | 95% CI | $p$ | $\beta$ | 95% CI | $p$ |
| Hippocampal-to-Ventricle Ratio | | | | | | |
| → g | **0.036** | **[0.015, 0.057]** | **9.00e-04** | **0.045** | **[0.022, 0.068]** | **1.55e-04** |
| → Speeds | 0.016 | [-0.013, 0.044] | 0.285 | 0.025 | [-0.009, 0.058] | 0.151 |
| Hippocampus (Residualized) | | | | | | |
| → g | **0.036** | **[0.017, 0.056]** | **2.10e-04** | 0.013 | [-0.008, 0.035] | 0.227 |
| → Speeds | 0.018 | [-0.009, 0.046] | 0.193 | **0.057** | **[0.025, 0.089]** | **4.04e-04** |
| Hippocampus (Unadjusted) | | | | | | |
| → g | **0.048** | **[0.026, 0.071]** | **2.36e-05** | **0.027** | **[0.002, 0.052]** | **0.035** |
| → Speeds | 0.025 | [-0.006, 0.056] | 0.119 | **0.068** | **[0.032, 0.104]** | **2.22e-04** |

Table S14: **SEM model fit indices.** Pooled models include sex as covariate; sex-stratified models estimate parameters separately. The HC (unadjusted) model uses a linear age term only; quadratic age was excluded due to multicollinearity between HC, ICV, and age$^2$. Good fit: CFI/TLI $> 0.95$, RMSEA $< 0.06$, SRMR $< 0.08$.

| Brain Measure | CFI | TLI | RMSEA | SRMR |
|---|---|---|---|---|
| Pooled | | | | |
| Hippocampal-to-Ventricle Ratio | 0.955 | 0.935 | 0.040 [0.039, 0.041] | 0.038 |
| Hippocampus (Residualized) | 0.953 | 0.932 | 0.040 [0.039, 0.041] | 0.037 |
| Hippocampus (Unadjusted) | 0.956 | 0.937 | 0.042 [0.041, 0.043] | 0.039 |
| Sex-Stratified | | | | |
| Hippocampal-to-Ventricle Ratio | 0.962 | 0.945 | 0.039 [0.037, 0.040] | 0.037 |
| Hippocampus (Residualized) | 0.959 | 0.941 | 0.039 [0.038, 0.040] | 0.037 |
| Hippocampus (Unadjusted) | 0.963 | 0.946 | 0.040 [0.039, 0.041] | 0.039 |

Table S15: **Factor loadings by sex.** $\lambda$ = standardized loading; SE = standard error. Bold indicates p < .05. Loadings constrained to 1.0 for scale identification not shown. Memory-Specific factor showed estimation failure in females and non-significant loadings in males (factor collapse); see Limitations.

| | Females | | | Males | | |
|---|---|---|---|---|---|---|
| Indicator | $\lambda$ | SE | $p$ | $\lambda$ | SE | $p$ |
| General Cognition ($g$) | | | | | | |
| Fluid Intelligence | **0.666** | **0.008** | **< 2.2e-16** | **0.673** | **0.008** | **< 2.2e-16** |
| Matrix Pattern Completion | **0.656** | **0.008** | **< 2.2e-16** | **0.630** | **0.010** | **< 2.2e-16** |
| Numeric Memory | **0.427** | **0.013** | **< 2.2e-16** | **0.427** | **0.066** | **7.25e-11** |
| Pairs Matching: Errors | **0.252** | **0.042** | **1.89e-09** | 0.253 | 0.544 | 0.642 |
| Pairs Matching: Time | **0.163** | **0.004** | **< 2.2e-16** | **0.176** | **0.005** | **< 2.2e-16** |
| Prospective Memory | **0.309** | **0.011** | **< 2.2e-16** | **0.336** | **0.011** | **< 2.2e-16** |
| Reaction Time | **0.013** | **0.000** | **< 2.2e-16** | **0.013** | **0.000** | **< 2.2e-16** |
| Symbol-Digit: Attempted | **0.394** | **0.016** | **< 2.2e-16** | **0.441** | **0.016** | **< 2.2e-16** |
| Symbol-Digit: Correct | **0.409** | **0.011** | **< 2.2e-16** | **0.463** | **0.013** | **< 2.2e-16** |
| Tower Rearranging | **0.532** | **0.010** | **< 2.2e-16** | **0.539** | **0.010** | **< 2.2e-16** |
| Trail Making: Alphanumeric | **0.460** | **0.014** | **< 2.2e-16** | **0.542** | **0.015** | **< 2.2e-16** |
| Trail Making: Numeric | **0.309** | **0.013** | **< 2.2e-16** | **0.324** | **0.019** | **< 2.2e-16** |
| Memory-Specific | | | | | | |
| Numeric Memory | — | — | — | -0.086 | 1.695 | 0.959 |
| Pairs Matching: Errors | — | — | — | 0.842 | 16.486 | 0.959 |
| Pairs Matching: Time | — | — | — | 1.135 | 22.929 | 0.961 |
| Prospective Memory | — | — | — | 0.003 | 0.152 | 0.984 |
| Processing Speed-Specific | | | | | | |
| Reaction Time | **0.431** | **0.012** | **< 2.2e-16** | **0.406** | **0.014** | **< 2.2e-16** |
| Symbol-Digit: Attempted | **0.454** | **0.015** | **< 2.2e-16** | **0.463** | **0.018** | **< 2.2e-16** |
| Symbol-Digit: Correct | **0.407** | **0.014** | **< 2.2e-16** | **0.425** | **0.016** | **< 2.2e-16** |
| Trail Making: Numeric | **0.224** | **0.017** | **< 2.2e-16** | **0.264** | **0.019** | **< 2.2e-16** |

Table S16: **SEM covariate effects on cognition.** $\beta$ = standardized coefficient; SE = standard error. Bold indicates p < .05. Memory-Specific factor not shown due to factor collapse.

| | Females | | | Males | | |
|---|---|---|---|---|---|---|
| Covariate | $\beta$ | SE | $p$ | $\beta$ | SE | $p$ |
| Hippocampal-Ventricle Ratio | | | | | | |
| Age | **-0.323** | **0.008** | **< 2.2e-16** | **-0.480** | **0.007** | **< 2.2e-16** |
| Age$^2$ | **-0.152** | **0.008** | **< 2.2e-16** | **-0.122** | **0.008** | **< 2.2e-16** |
| Deprivation Index | 0.003 | 0.008 | 0.682 | -0.008 | 0.008 | 0.360 |
| Education | **-0.027** | **0.008** | **0.002** | **-0.021** | **0.009** | **0.019** |
| Intracranial Volume | **-0.137** | **0.008** | **< 2.2e-16** | **-0.124** | **0.009** | **< 2.2e-16** |
| General Cognition ($g$) | | | | | | |
| Age | **-0.200** | **0.012** | **< 2.2e-16** | **-0.249** | **0.013** | **< 2.2e-16** |
| Age$^2$ | **-0.085** | **0.010** | **< 2.2e-16** | **-0.076** | **0.011** | **4.42e-12** |
| Deprivation Index | **-0.057** | **0.011** | **8.96e-08** | **-0.082** | **0.012** | **2.07e-12** |
| Education | **0.347** | **0.010** | **< 2.2e-16** | **0.363** | **0.011** | **< 2.2e-16** |
| Intracranial Volume | **0.174** | **0.010** | **< 2.2e-16** | **0.154** | **0.011** | **< 2.2e-16** |
| Processing Speed-Specific | | | | | | |
| Age | **-0.703** | **0.016** | **< 2.2e-16** | **-0.676** | **0.020** | **< 2.2e-16** |
| Age$^2$ | **-0.055** | **0.016** | **4.93e-04** | -0.033 | 0.017 | 0.056 |
| Deprivation Index | **-0.050** | **0.014** | **3.88e-04** | **-0.071** | **0.017** | **2.00e-05** |
| Education | **-0.038** | **0.018** | **0.033** | **0.045** | **0.019** | **0.016** |
| Intracranial Volume | 0.016 | 0.015 | 0.285 | -0.001 | 0.016 | 0.929 |

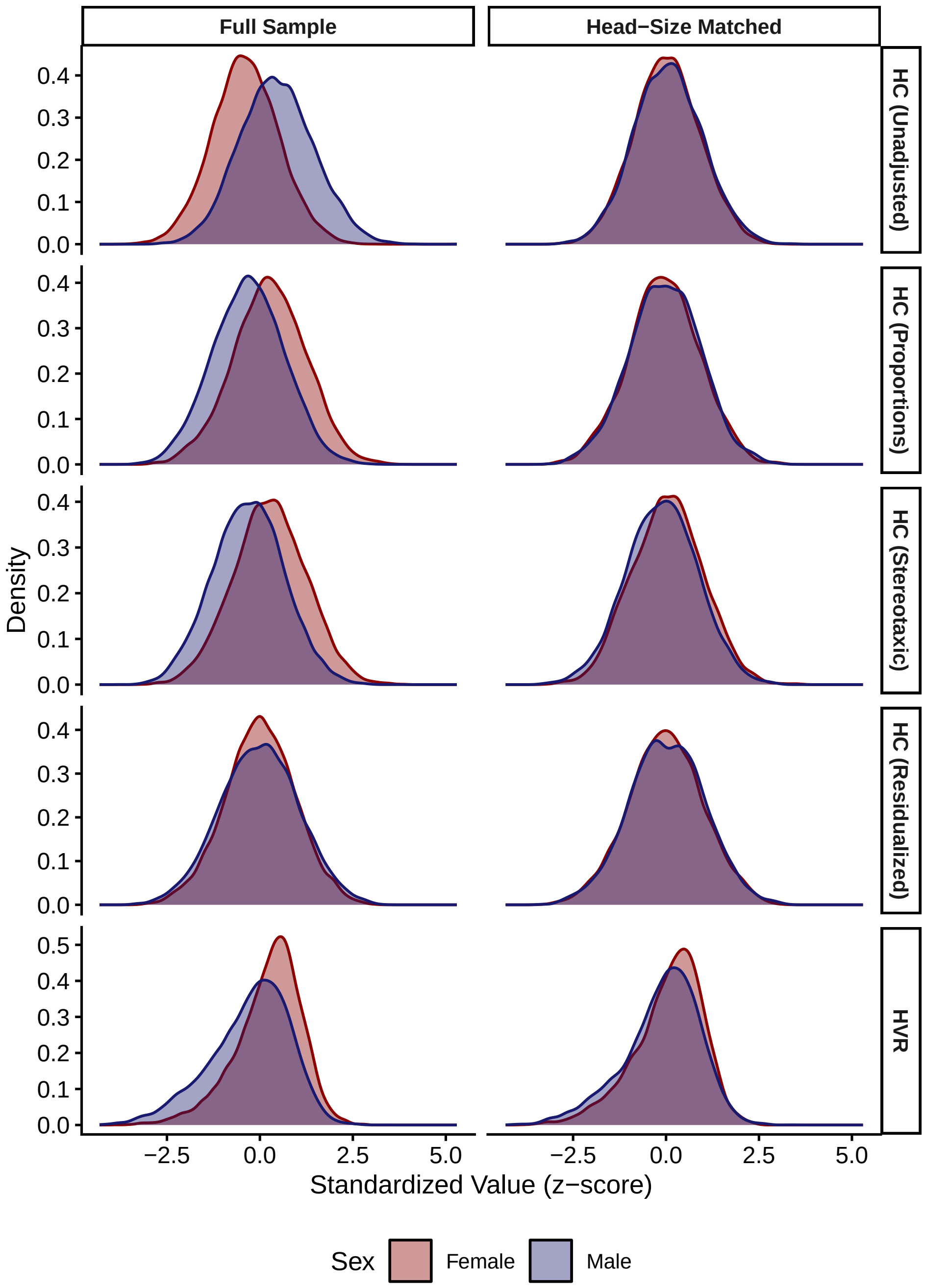


Figure S2: **Distribution overlap by sex.** Across adjustment methods. Left column: full sample. Right column: head-size matched sample (age ±1 year, intracranial volume ±25 cc).

### Appendix D: Sensitivity Analyses

Table S17: **Site-adjusted sex difference effect sizes.** ICC = intraclass correlation coefficient indicating proportion of variance attributable to site.

| | | Effect Size | |
|---|---|---|---|
| Adjustment | Site ICC | d | 95% CI |
| Hippocampus | | | |
| Unadjusted | 0.012 | -0.885 | [-0.908, -0.861] |
| Proportions | 0.004 | 0.584 | [0.560, 0.607] |
| Residuals | 0.006 | 0.025 | [0.001, 0.049] |
| Stereotaxic | 0.007 | 0.593 | [0.569, 0.616] |
| Lateral Ventricles | | | |
| Unadjusted | 0.012 | -0.843 | [-0.866, -0.819] |
| Proportions | 0.010 | -0.343 | [-0.367, -0.319] |
| Residuals | 0.009 | -0.163 | [-0.187, -0.139] |
| Stereotaxic | 0.010 | -0.309 | [-0.333, -0.285] |
| Hippocampus-to-Ventricle ratio | | | |
| Self-normalizing | 0.011 | 0.520 | [0.497, 0.544] |

Table S18: **Sex differences by hemisphere.** HC/LV = ICV-residualized; HVR = self-normalizing ratio. d = Cohen's d (positive = females > males).

| Hemisphere | d | 95% CI |
|---|---|---|
| Hippocampal-to-Ventricle Ratio | | |
| Left | -0.007 | [-0.031, 0.017] |
| Right | 0.050 | [0.026, 0.074] |
| Hippocampus | | |
| Left | -0.143 | [-0.167, -0.120] |
| Right | -0.160 | [-0.183, -0.136] |
| Lateral Ventricles | | |
| Left | 0.477 | [0.453, 0.502] |
| Right | 0.481 | [0.457, 0.505] |

Table S19: **Sensitivity to psychiatric exclusions.** Sensitivity sample includes participants with ICD-10 F00-F99 diagnoses. Δd = difference in Cohen's d; values near zero indicate robust findings.

| | Primary Sample | | | Sensitivity Sample | | | |
|---|---|---|---|---|---|---|---|
| Adjustment | N | d | 95% CI | N | d | 95% CI | Δd |
| Hippocampus | | | | | | | |
| Unadjusted | 27,680 | -0.885 | [-0.910, -0.860] | 30,032 | -0.886 | [-0.910, -0.862] | -0.001 |
| Proportions | 27,680 | 0.580 | [0.556, 0.605] | 30,032 | 0.571 | [0.548, 0.595] | -0.009 |
| Residuals | 27,680 | 0.022 | [-0.002, 0.046] | 30,032 | 0.016 | [-0.006, 0.039] | -0.006 |
| Stereotaxic | 27,680 | 0.588 | [0.563, 0.612] | 30,032 | 0.580 | [0.556, 0.603] | -0.008 |
| Lateral Ventricles | | | | | | | |
| Unadjusted | 27,680 | -0.840 | [-0.865, -0.815] | 30,032 | -0.841 | [-0.865, -0.818] | -0.002 |
| Proportions | 27,680 | -0.341 | [-0.365, -0.317] | 30,032 | -0.345 | [-0.368, -0.322] | -0.004 |
| Residuals | 27,680 | -0.162 | [-0.185, -0.138] | 30,032 | -0.165 | [-0.188, -0.142] | -0.003 |
| Stereotaxic | 27,680 | -0.308 | [-0.332, -0.284] | 30,032 | -0.312 | [-0.335, -0.289] | -0.004 |
| Hippocampus-to-Ventricle ratio | | | | | | | |
| HVR (Self-Normalizing) | 27,680 | 0.517 | [0.493, 0.542] | 30,032 | 0.519 | [0.495, 0.542] | 0.001 |

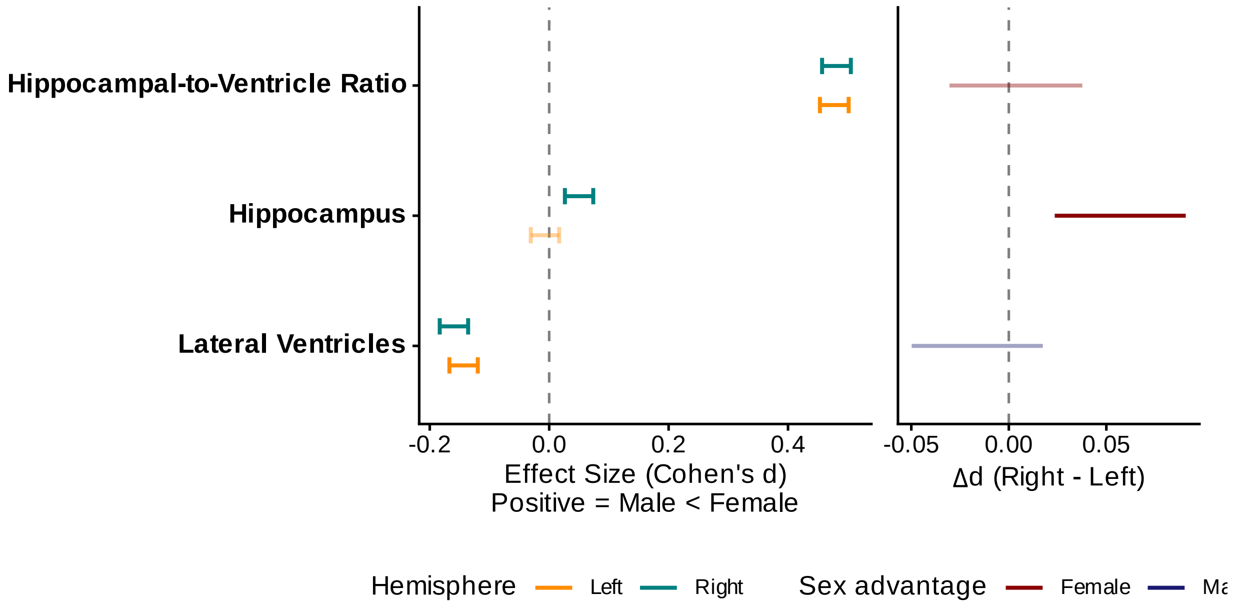


Figure S3: **Hemisphere comparison of sex differences.** Faded error bars indicate non-significant effects. Right panel shows the right minus left difference with 95% confidence interval, colored by overall sex advantage direction (red = female advantage, blue = male advantage; faded = non-significant difference).

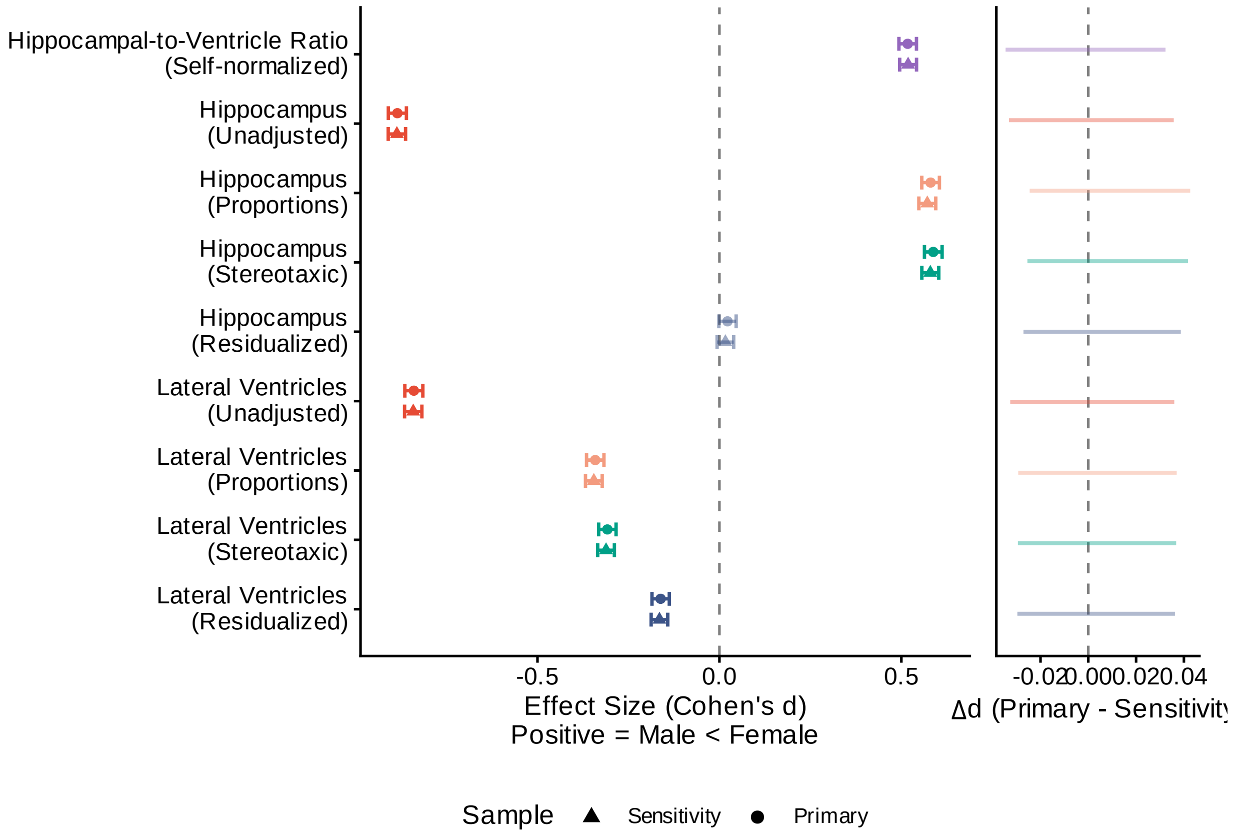


Figure S4: **Sensitivity analysis for psychiatric exclusions.** Faded points indicate non-significant effects. Right panel shows the primary minus sensitivity difference with 95% confidence interval.
